# Using maximum entropy production to describe microbial biogeochemistry over time and space in a meromictic pond

**DOI:** 10.1101/266346

**Authors:** Joseph J. Vallino, Julie A. Huber

## Abstract

The maximum entropy production (MEP) conjecture posits that systems with many degrees of freedom will likely organize to maximize the rate of free energy dissipation. Previous work indicates that biological systems can outcompete abiotic systems by maximizing free energy dissipation over time by utilizing temporal strategies acquired and refined by evolution, and over space via cooperation. In this study, we develop an MEP model to describe biogeochemistry observed in Siders Pond, a phosphate limited meromictic system located in Falmouth, MA that exhibits steep chemical gradients due to density-driven stratification that supports anaerobic photosynthesis as well as microbial communities that catalyze redox cycles involving O, N, S, Fe and Mn. The MEP model uses a metabolic network to represent microbial redox reactions, where biomass allocation and reaction rates are determined by solving an optimization problem that maximizes entropy production over time and a 1D vertical profile constrained by an advection-dispersion-reaction model. We introduce a new approach for modeling phototrophy and explicitly represent aerobic photoautotrophs, anoxygenic photoheterotrophs and anaerobic photoautotrophs. The metabolic network also includes reactions for heterotrophic bacteria, sulfate reducing bacteria, sulfide oxidizing bacteria and aerobic and anaerobic grazers. Model results were compared to observations of biogeochemical constituents collected over a 24 hour period at 8 depths at a single 15 m deep station in Siders Pond. Maximizing entropy production over long (3 d) intervals produced results more similar to field observations than short (0.25 d) interval optimizations, which support the importance of temporal strategies for maximizing entropy production over time. Furthermore, we found that entropy production must be maximized locally instead of globally where energy potentials are degraded quickly by abiotic processes, such as light absorption by water. This combination of field observations with modeling results show that microbial systems in nature can be accurately described by the maximum entropy production conjecture applied over time and space.

## 1 Introduction

Mass and energy flow associated with the growth and predation of bacteria, archaea and eucaryotes in microbial food webs coupled with abiotic reactions and transport processes define biogeochemical cycles that occur in ecosystems ranging in size from less than a liter (Marino et al., 2016) to the entire planet. Because organisms are ultimately responsible for most observed biogeochemical transformations, it is customary and natural to focus on the bioenergetics and growth and predation of the organisms that constitute food webs in order to understand and predict biogeochemical transformations. This organismal focus has a long history and has advanced our understanding and prediction of ecosystem dynamics and the mass and energy flow through them (Riley, 1946; Fasham et al., 1990; Le Quere et al., 2005; Friedrichs et al., 2007; Schartau et al., 2017). While focusing on the growth, predator-prey and cooperative interactions of organisms will continue to contribute to our understanding of ecosystem bioenergetics, there are some challenges that limit this approach for microbial systems that form the basis of biogeochemical cycles. Microbial communities are diverse, complex and abundant, consisting, for example, of upwards of 10^9^ microorganisms per liter of seawater, with estimates that Earth hosts close to 1 trillion microbial species (Sogin et al., 2006; Locey and Lennon, 2016). With the advent of metagenomics, next generation sequencing and bioinformatics, the challenging task of deciphering and annotating the metabolic capabilities and activities of bacteria and other microorganisms has begun, but untangling the Gordian knot will take significantly more time and investment (Ponomarova and Patil, 2015), and determining how information in genomes contributes to competition and cooperation is still in its infancy (Hallam and McCutcheon, 2015; Pasternak et al., 2015; Worden et al., 2015). Even more challenging is understanding and predicting community composition dynamics and succession as environmental conditions change, both from exogenous and endogenous drivers (Konopka et al., 2015). To develop predictive models of biogeochemistry based on organisms and the genes they carry, exchange horizontally and differentially express requires an enormous amount of information, which will likely take many decades to compile and decipher, but good progress is being made (Reed et al., 2014; Coles et al., 2017). While we believe this reductionist approach is essential, there is also a complementary approach to understanding microbial biogeochemistry that is less studied and uses a more thermodynamic, or whole systems, approach.

Understanding how ecosystems function at the systems level has a long tradition in theoretical ecology (Chapman et al., 2016; Vallino and Algar, 2016), but the underlying premise is that ecosystems organize so as to maximize an objective function, such as maximizing power proposed by Lotka (1922) nearly 100 years ago. The advantage of the systems approach is that the variational principle can be used to determine how an ecosystem will organize and function without the knowledge of which organisms are present and how their population changes over time, provided there is sufficient diversity. That is, understanding and modeling of ecosystems can focus on function rather than on organisms, and there is growing support that stable function can arise from dynamic communities (Fernandez et al., 1999; Fernandez-Gonzalez et al., 2016; Louca and Doebeli, 2016; Needham and Fuhrman, 2016; Coles et al., 2017). Here, we build upon the conjecture that microbial systems organize to maximize entropy production. The maximum entropy production (MEP) principle proposes that systems (biotic systems included) with sufficient degrees of freedom will likely organize to maximize the dissipation of Gibbs free energy (Dewar, 2003; Lorenz, 2003; Martyushev and Seleznev, 2006). The MEP principle has been applied to both abiotic and biotic processes (Kleidon and Lorenz, 2005; Kleidon et al., 2010; Dewar et al., 2014), and we have used MEP to model periodically forced methanotrophic microbial communities (Vallino et al., 2014) and investigate metabolic switching in nitrate reducing environments (Algar and Vallino, 2014).

An unresolved aspect of MEP involves the temporal and spatial scales over which it operates. MEP theory has only been derived for nonequilibrium steady-state systems, but microbial systems are dynamic, so steady state does not apply. Consequently, we have proposed that abiotic processes, such as fire or a rock rolling down a hill, maximize instantaneous entropy production; that is, they follow a steepest descent trajectory down a potential energy surface. Biological systems, however, use information acquired by evolution and culled by natural selection to follow alternate pathways that maximize entropy production over time, which allows biology to sometimes outcompete abiotic processes (Vallino, 2010). Similarly, when considering a spatial domain, entropy production can either be maximized locally at each point in the domain or entropy production at each point can be adjusted so that entropy production is maximized globally over the entire domain. A simple numerical study indicated that when a system organizes over space, entropy produced by global optimization can exceed that from local optimization, but this requires spatial coordination by the community, while abiotic systems are likely to only maximize entropy production locally (Vallino, 2011). This paper seeks to examine both of these ideas using spatially and temporally distinct observations collected from a field site.

In this study we develop an MEP-based model to predict microbial biogeochemistry in a meromictic pond located in Falmouth, MA (Siders Pond) that includes metabolic processes for phytoplankton, green sulfur bacteria, heterotrophic bacteria, sulfate reducing bacteria, sulfide oxidizing bacteria, photoheterotrophs and aerobic and anaerobic predators, but the primary objective of the model is to dissipate energy potentials, not grow organisms. While previous MEP models have been developed for chemolithoautotrophs, chemolithoheterotrophs and chemoorganoheterotrophs, this study expands the metabolic reaction repertoire to include photoautotrophs and photoheterotrophs. The approach is also extended to include an explicit spatial dimension, and we compare model output to observations in Siders Pond collected over a 24 hour sampling period from eight depths. MEP solutions using two different optimization timescales (0.25 d versus 3.0 d) are contrasted and compared to observations, and we discuss the problem of local versus global MEP optimization for energy potentials that are quickly dissipated abiotically, such as light. Our results suggest that microbial systems in nature can be accurately described by the maximum entropy production conjecture applied over time and space.

## 2 Methods

This section describes the development of the MEP model, including the 1D transport model for Siders Pond, followed by Siders Pond sample collection procedures and analytical methods used for sample analyses.

### 2.1 Model Development

The equations used to model biogeochemistry in Siders Pond are provided in detail in the *Supplementary Material (SM)*, so the descriptions in this section focus primarily on model concepts that form the basis of the modeling approach and extensions to previous studies. The model consists of three main components: 1) a 1D transport model; 2) a set of biologically catalyzed reactions that constitute the metabolic network of the microbial community, including predators such as protist and viruses; 3) an optimization component in which control variables that govern reaction stoichiometry, kinetics and thermodynamics are determined so as to maximize internal entropy production over a specified interval of time and space. One of the underlying objectives of the approach is to replace what are typically poorly constrained biological parameters, such as maximum specific growth rates, substrate affinities, prey preferences, reaction efficiencies, etc, with optimal control variables whose values are determined by maximizing free energy dissipation. Consequently, these MEP-based models contain relatively few adjustable parameters, but see Section 2.1.3.1 below. We describe briefly below the three model components, but we focus first on the representation of the metabolic network, as this forms the foundation of the approach, and includes the concentrations of 11 chemical constituents and 8 functional groups (Fig. 1): salt, 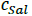; dissolved oxygen, 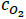; dissolved inorganic carbon, 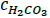; inorganic phosphate, 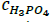; sulfate, 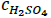; hydrogen sulfide, 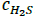; phytoplankton carbohydrates, 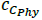; green sulfur bacteria carbohydrates, 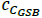; labile organic carbon, 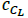; refractory organic carbon, 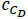; refractory organic phosphate, 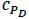; phytoplankton, 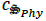; green sulfur bacteria, 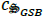; aerobic predators, 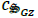; anaerobic predators, 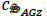; heterotrophic bacteria, 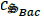; sulfate reducing bacteria, 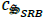; photoheterotrophs, 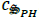; sulfide oxidizing bacteria, 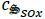.

**Fig. 1.**
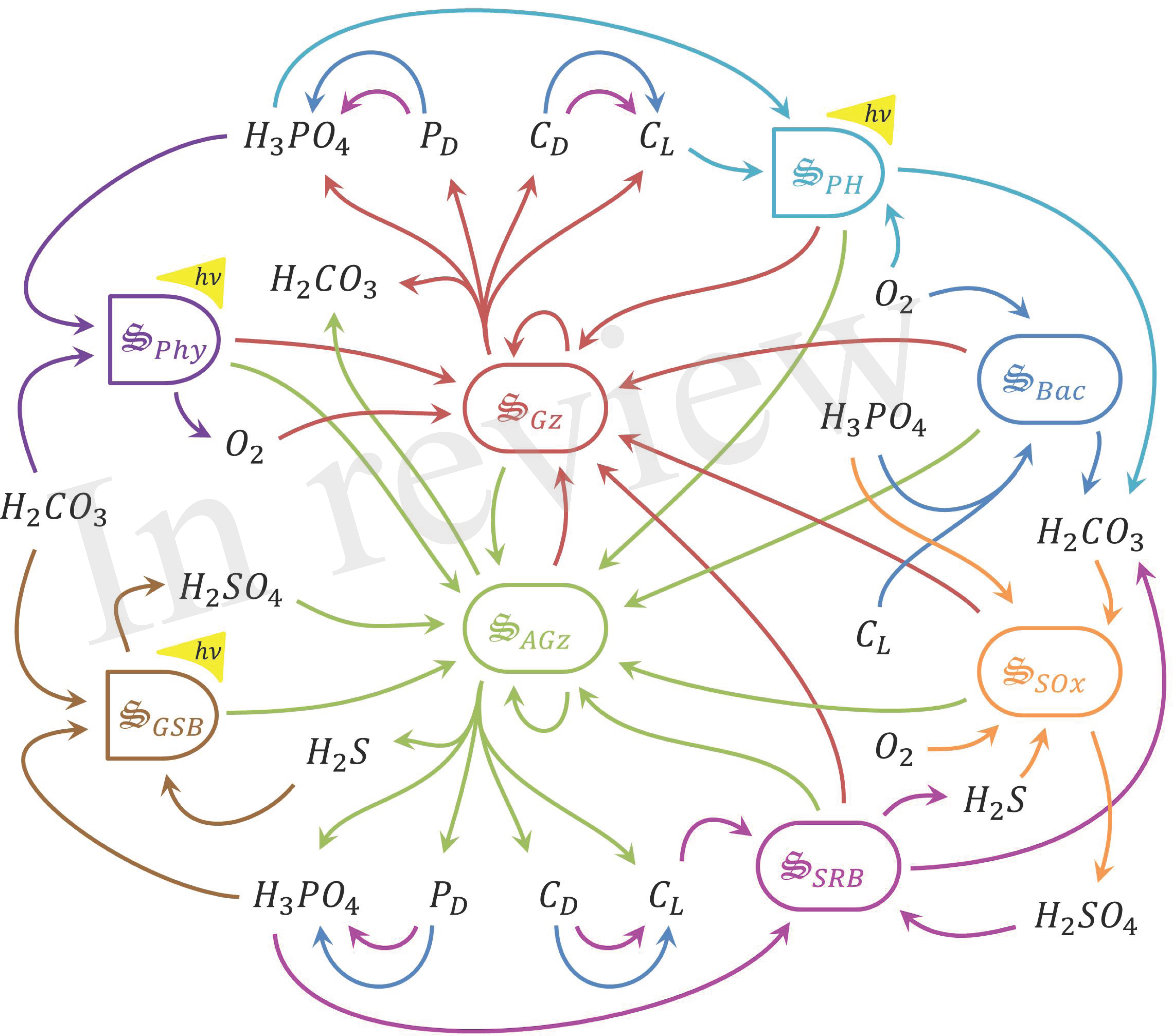
Schematic of catalysts and associated reactions used in the MEP model for Siders Pond. Functional groups include: 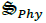, phytoplankton, purple, 2 rxns; 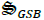, green sulfur bacteria, brown, 2 rxns; 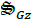, aerobic grazers, red, 8 rxns; 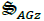, anaerobic grazers, green, 8 rxns; 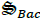, heterotrophic bacteria, blue, 3 rxns; 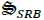, sulfate reducing bacteria, magenta, 3 rxns; 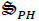, photoheterotrophs, cyan, 1 rxn; 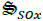, sulfide oxidizing bacteria, orange, 1 rxn. Other abbreviations: *hv*, photon capture; C_*L*_, labile organic carbon; C_*D*_, refractory organic carbon; P_*D*_, refractory organic phosphorous. See Table 1 for qualitative representation of functional reactions and Section 2.7 of the *Supplementary Material* for stoichiometrically balanced reactions.

**Table 1.** Reactions associated with the 8 biological catalysts, 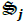, used to model microbial biogeochemistry in Siders Pond, where ***r_i,j_*** represents sub-reaction ***i*** of biological catalyst 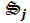. There are a total of 28 reactions, where 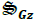 and 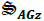 each catalyzed 8 sub-reactions. Reactions are shown to emphasize function only. Complete reaction stoichiometries, including influence of optimal control variables, are given in Section 2.7 of the *Supplementary Material*. See caption of Figure 1 for nomenclature.

| Rxn. | Abbreviated Stoichiometry | Cat. |
| --- | --- | --- |
| $r_{1,Phy}$ | $H_2CO_3 + h\nu \rightarrow C_{Phy} + O_2$ | $\mathfrak{S}_{Phy}$ |
| $r_{2,Phy}$ | $C_{Phy} + NH_3 + H_3PO_4 + O_2 \rightarrow \mathfrak{S}_{Phy} + H_2CO_3$ | $\mathfrak{S}_{Phy}$ |
| $r_{1,GSB}$ | $H_2CO_3 + H_2S + h\nu \rightarrow C_{GSB} + H_2SO_4$ | $\mathfrak{S}_{GSB}$ |
| $r_{2,GSB}$ | $C_{GSB} + NH_3 + H_3PO_4 + H_2SO_4 \rightarrow \mathfrak{S}_{GSB} + H_2CO_3 + H_2S$ | $\mathfrak{S}_{GSB}$ |
| $r_{1-8,GZ}$ | $\mathfrak{S}_i + C_i + O_2 \rightarrow \mathfrak{S}_{GZ} + H_2CO_3 + C_D + NH_3 + N_D + H_3PO_4 + P_D + C_L$ | $\mathfrak{S}_{GZ}$ |
| $r_{1-8,AGZ}$ | $\mathfrak{S}_i + C_i + H_2SO_4 \rightarrow \mathfrak{S}_{AGZ} + H_2CO_3 + C_D + NH_3 + N_D + H_3PO_4 + P_D + H_2S + C_L$ | $\mathfrak{S}_{AGZ}$ |
| $r_{1,Bac}$ | $C_L + NH_3 + H_3PO_4 + O_2 \rightarrow \mathfrak{S}_{Bac} + H_2CO_3$ | $\mathfrak{S}_{Bac}$ |
| $r_{2,Bac}$ | $C_D \rightarrow C_L$ | $\mathfrak{S}_{Bac}$ |
| $r_{3,Bac}$ | $P_D \rightarrow H_3PO_4$ | $\mathfrak{S}_{Bac}$ |
| $r_{1,SRB}$ | $C_L + NH_3 + H_3PO_4 + H_2SO_4 \rightarrow \mathfrak{S}_{SRB} + H_2CO_3 + H_2S$ | $\mathfrak{S}_{SRB}$ |
| $r_{2,SRB}$ | $C_D \rightarrow C_L$ | $\mathfrak{S}_{SRB}$ |
| $r_{3,SRB}$ | $P_D \rightarrow H_3PO_4$ | $\mathfrak{S}_{SRB}$ |
| $r_{1,PH}$ | $C_L + NH_3 + H_3PO_4 + h\nu \rightarrow \mathfrak{S}_{PH} + H_2CO_3$ | $\mathfrak{S}_{PH}$ |
| $r_{1,SOx}$ | $H_2CO_3 + H_2S + O_2 + NH_3 + H_3PO_4 \rightarrow \mathfrak{S}_{SOx} + H_2SO_4$ | $\mathfrak{S}_{SOx}$ |

#### 2.1.1 Catalysts and metabolic reaction network

Our approach views a complex microbial community as a collection of catalysts (denoted with the special symbol 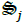) that each have a subset of *n_r,j_* functional reactions they catalyze. The catalyst has an elemental composition given by 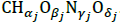. For this study all catalysts are assigned the same composition as yeast with associated thermodynamic properties (Battley et al., 1997), but this is not an overly constraining approximation (Vallino et al., 2014) and can be easily relaxed if needed. The catalysts and reactions included in the metabolic network represent the capabilities of the entire microbial community, but the reactions are distributed across 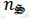 catalysts (8 for the case of Siders Pond, Fig. 1), just as functions are distributed across phyla in natural communities (Vallino, 2003). The reactions are highly simplified and condensed, and consist of two essential components: an anabolic reaction that synthesizes catalyst from environmental resources, and a catabolic reaction that provides free energy to drive the anabolic reaction forward. A highly simplified list of metabolic reactions for the Siders Pond model is given in Table 1. To convey the basic ideas, we consider below the phylum responsible for sulfate reduction, but see the *SM* for the complete derivation.

The catalyst 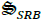 represents the capabilities of sulfate reducing bacteria (SRB) that oxidize labile organic matter, C_*L*_, with an elemental composition CH_2_O, using sulfate as the electron acceptor. The anabolic and catabolic reactions are given by,

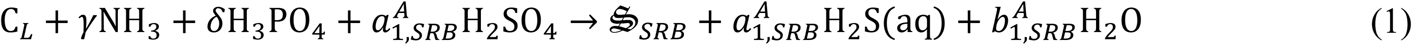

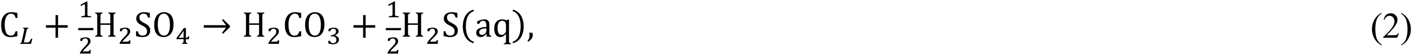

respectively, where the stoichiometric coefficients, 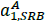 and 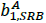, are determined from elemental balances around O and H, since C, N and P are in balance. An important concept for the approach is that both reactions above must be catalyzed by 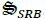, so the reaction rate is proportional to the concentration of the catalyst, 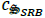 or 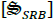, present. These two reactions can be combined by introducing a reaction weighting factor, or growth efficiency variable, *ε_SRB_*, to produce an overall reaction representing sulfate reducing bacteria, *r*_1,*SRB*_, given by,

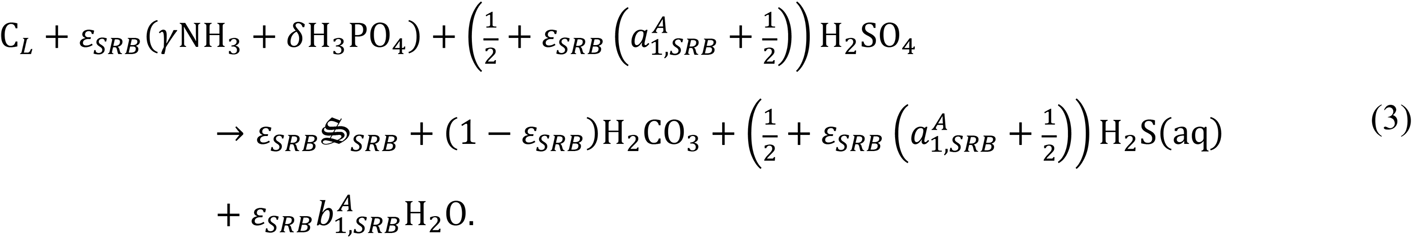

The reaction efficiency variable, *ε_SRB_*, is ***one of two classes of optimal control variables*** and a central design feature of the MEP model. As *ε_SRB_* approaches 1, Eq. (3) represents 100# conversion of labile carbon plus N and P resources to catalyst, while as *ε_SRB_* approaches 0, the reaction changes to 100% anaerobic combustion of labile carbon. From an entropy production perspective, only the catabolic reaction dissipates significant free energy, so *ε_SRB_* should be set to zero to maximize entropy production; however, the catabolic reaction cannot proceed without catalyst. There exists, then, an optimum value of *ε_SRB_* that produces just enough 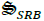 catalyst to dissipate the chemical potential between C_*L*_ and H_2_SO_4_, but *ε_SRB_* must change as a function of C_*L*_ and H_2_SO_4_ supply rates as well as N and P availability. Conceptually, the MEP problem is to determine how *ε_SRB_* should change over time and space to maximize free energy dissipation; however, there are a few other important details to fill in.

Since we have discussed MEP model development in previous work (Vallino, 2010; Vallino, 2011; Algar and Vallino, 2014; Vallino et al., 2014) and have provided the model equations in the *Supplementary Material*, we only briefly discuss the main ideas followed by modifications and new additions for phototrophs. As before, we use Alberty’s (2003) approach to calculate Gibbs free energy of reaction, 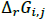, as a function of *ε_j_*, temperature, *T*, and pH, which accounts for dissociation of chemical species (i.e., 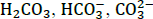, etc), and ionic strength is used to approximate activities from reactant and product concentrations. For non-phototrophic reactions (i.e., chemotrophs), an adaptive Monod equation parameterized by *ε_j_* accounts for the tradeoffs between substrate affinity, maximum specific growth rate and growth efficiency and can approximate oligotrophic to copiotrophic growth kinetics (such as SAR11 growing in the Sargasso Sea to *E. coli* growing on LB medium in the lab) as *ε_j_* varies between 0 and 1 (Algar and Vallino, 2014; Vallino et al., 2014). In addition to the kinetic constraints, *F_K_*, reaction rates, *r_i,j_*, are also constrained by reaction thermodynamics, *F_T_*, as described by La Rowe et al. (2012) (see Eq. S24). Consequently, the rates of chemical reactions (see Section 2.7.6 of *SM*) take the following general form,

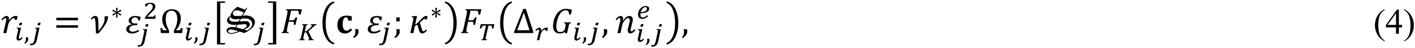

where vector **c** is the concentration of substrates and products, 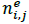 is the number of electrons transferred in the redox reaction, 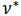 and 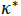 are universal constants (we have used the same values of 350 d^-1^ for 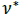 and 5000 mmol m^-3^ for 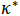 for all models to date) and Ω_*i,j*_ is the ***second optimal control variable class***. Since a catalyst, 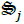, can catalyze *n_r,j_* sub-reactions, Ω_*i,j*_ determines how much of catalyst 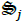 is allocated to reaction *r_i,j_* at a given time and location in space. For instance, the sulfate reducing bacteria are also given two reactions in addition to Eq. (3) (Table 1). Reactions *r_2,SRB_* and *r_3,SRB_* allow SRB to decompose recalcitrant organic carbon, C_*L*_, and phosphorous, P_*D*_, into labile organic carbon, C_*L*_, and inorganic phosphate, respectively, but to do so they must deallocate from biosynthesis reaction, *r_1,SRB_*, Eq. (3). That is, each ε_*i,j*_ is bound between 0 and 1, and Ω_*i,j*_ must sum to unity over *i* for each 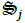. An example of how 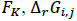, and *F_T_* vary over time and space to influence reaction rate is given in *SM* Section 4.1, Fig. S3 for growth of sulfate reducing bacteria, *r_1,SRB_*.

Entropy production per unit volume for chemotrophic reactions, 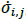, is given by,

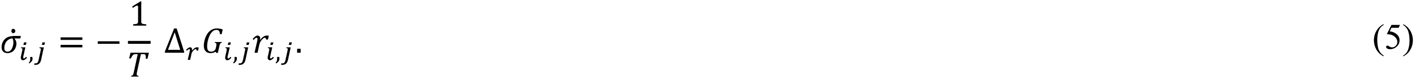

In the formulation above, the free energy of reaction, 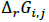, for the pure anabolic reaction, Eq. (1), might be positive (but it can be negative, see Vallino, 2010). To improve computational consistency between anabolic reactions with differing free energies, the reaction stoichiometries are augmented to insure that 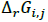 goes to zero as goes to 1 for the anabolic component of the reaction. For an anabolic reaction that has a strongly positive 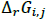, the reaction is coupled to the catabolic reaction based on the ratio of the anabolic to catabolic reaction free energies, defined as *n_i,j_* in the stoichiometric equations. Consequently the true stoichiometry for *r_1,SRB_*, as well as other reactions in the network, is slightly more cluttered (e.g., Eq. S103), but still readily calculated.

##### 2.1.1.1 Phototrophy

In this version of the MEP model we introduce catalysts associated with phototrophic growth, specifically phytoplankton (Phy), 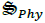, anaerobic green sulfur bacteria (GSB), 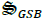, and photoheterotrophs (PH), 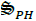. Both Phy and GSB are modeled similarly using two sub-reactions (Table 1). One sub-reaction couples photon capture that drives CO_2_ fixation into carbohydrates, C_*phy*_ and C_*GSB*_, with elemental composition CH_2_O, and a second sub-reaction converts carbohydrates into biomass in a manner analogous to growth of heterotrophic bacteria and GSB previously developed. We focus on phytoplankton here (but see *SM* Section 2.7.2) because development for GSB is similar, and can be found in *SM* Section 2.7.3.

The carbon fixation reaction for Phy is given by,

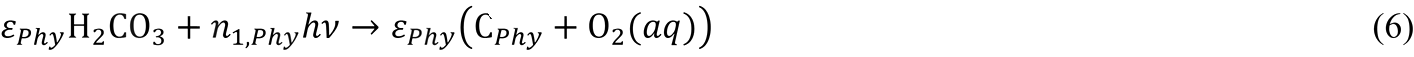

where *hv* are captured photons of frequency *v, h* is Plank’s constant and *n_1,Phy_* is the mmoles of photons needed to fix 1 mmol of CO_2_ at 100% efficiency (i.e., *ε_Phy_* = 1) (note, units for reaction rates in the model are in mmol m^-3^d^-1^, so we use mmol instead of mol in this description). The Gibbs free energy for a mmol of photons is given by

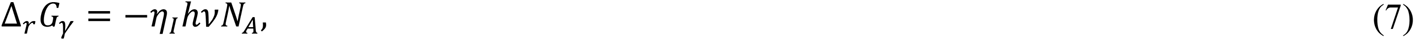

where *N_A_* is Avogadro’s number (in mmol) and *η_I_* is the thermodynamic efficiency for converting electromagnetic radiation into work (see Eq. (S26) and Candau, 2003). If we define 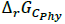 as the Gibbs free energy for fixing 1 mmol of H_2_CO_3_ into C_*phy*_ plus O_2_ (see Eq. (S29)), then

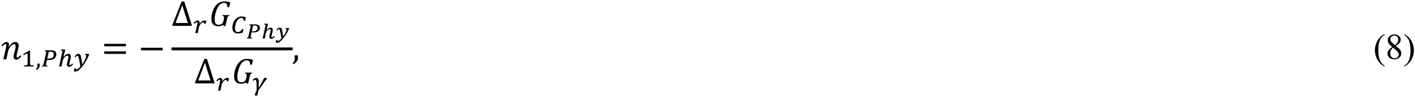

and the Gibbs free energy of reaction for Eq. (6) is given by,

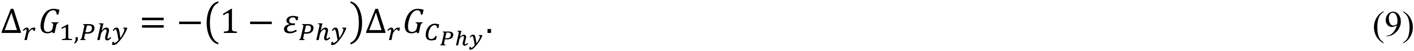

In this formulation, *ε_Phy_* governs the efficiency for the conversation of electromagnetic potential into chemical potential. If 100% of solar radiation is converted to chemical potential, no entropy is produced, and the free energy of reaction for Eq. (6) is zero, so the reaction will not proceed due to thermodynamic constraints. However, as *ε_Phy_* is reduced below 1, some photons in the photosynthetic active radiation (PAR) spectrum are simply dissipated as heat, and all are dissipated as heat when *ε_Phy_* =0. In Eq. (6), the classic quantum yield, *φ*, (Kishino et al., 1986) is given by 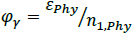, which obviously varies as a function of the optimal control variable, *ε_Phy_*.

The reaction rate for Eq. (6) depends on the rate of photon capture, which is given by,

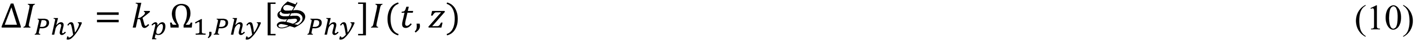

where *k_p_* is the coefficient for light absorption by particles, 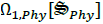 is the fraction of phytoplankton catalytic machinery allocated to capturing photons (i.e., chlorophyll, other pigments, electron transfer chains, etc), and *I*(*t,Z*) is the light intensity (mmol photons m^-2^d^-1^) at time *t* and depth *z* (see *SM* Section 2.5). Consequently, the reaction rate for Eq. (6) is given by,

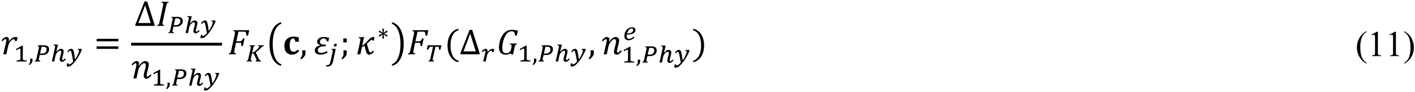

This reaction has similarities to those typically used to describe phytoplankton growth (Macedo and Duarte, 2006), but our derivation focuses not on the local light intensity level, but rather on how much light is actually intercept by the phytoplankton, as governed by 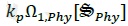, and how much of that free energy is actually used to drive carbon fixation, as governed by *ε_Phy_*. As evident in Eq. (11), the maximum rate is directly tied to the rate of photon interception, ∆*I_Phy_*, not by an arbitrary maximum specific growth parameter. The second reaction used by Phy and GSB (Table 1) is simply the conversion of reduced organic carbon, C_*Phy*_ and C_*GSB*_, into more catalyst, but of course C_*Phy*_ and C_*GSB*_ can also just be combusted to produce entropy depending of the value of *ε_Phy_*. The carbohydrate pools serve a similar function as cell quota in Droop’s (1973) model.

The reaction for photoheterotrophs (PH) differs slightly from that above (*SM* Section 2.7.8). In this case only one reaction is used (*r_1,PH_*, Table 1), where the photon capture is linked to the conversion of labile carbon into PH catalyst. As above, photons captured can also be dissipated as heat for *ε_PH_* < 1, or the free energy can be used to drive biosynthesis (*ε_PH_* > 1), and photon energy replaces catabolic reaction used in chemotroph reactions. In the currently formulation, reaction rate is also tied to photon capture rate, so growth can only proceed during the day (Eq. (S131)). A second chemotrophic sub-reaction (like *r_1,Bac_*, Eq. (S93)) could have been added to 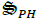, but this would increase the number of optimal control variables in the problem, and would be redundant with 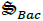, so this option was not explored.

#### 2.1.2 Entropy production

It is outside the scope of this manuscript to delve into the thickets of entropy (see Vallino and Algar, 2016), but a few definitions are in order, since entropy accounting is the foundation of the MEP approach. Simply stated, entropy production occurs when an energy potential is destroyed and dissipated as heat to the environment, but not when the potential is converted from one potential to another, if done so reversibly, which technically is possible, but never occurs in practice. For example, a flame converting chemical potential into heat or light being absorbed by water both result in maximum entropy production; these are irreversible processes. On the other hand, entropy is not produced by converting electromagnetic potential into chemical potential, which phytoplankton do partially, but they also convert a fraction of that potential to heat, as dictated by *ε_Phy_* in the model. As *ε_Phy_* approaches 1, electromagnetic potential is converted to chemical potential without loss, but *F_T_* (Eq. S24) drivers the reaction rate to 0. For organisms to grow at a non-zero rate an energy potential must be partially dissipated and some entropy must be produced.

For the Siders Pond model, entropy production associated electromagnetic radiation can be produced along three pathways. Photons intercepted by water, *k_w_I*(*t,Z*), and particles lacking photochemical properties, such as bacteria, 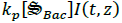 and grazers, 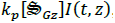, as well as by the non-photosynthetic components of phytoplankton, 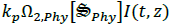 and 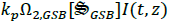, are simply dissipated as heat (long wave radiation), and the entropy produced from them is readily calculated. Photons intercepted by the photosynthetic machinery of phototrophs can be either dissipated as heat, *ε_j_* = 0, or the energy potential can be transferred to chemical potential, *ε_j_* = 1, but in general it is a combination of those processes,0 < *ε_j_* < 1. This latter route is considered entropy production by reaction in the Results Section, even though it involves photons. Consequently, total entropy production, 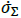 (Eq. (S178) first expression) is the sum of three processes: entropy production by photon absorption by water, 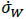 (Eq. (S176)), entropy production by light absorption by particles, 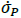 (Eq. (S177)) and entropy production by reaction, 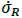 (Eq. (S178) second expression). In the photic zone of Sider Pond, it is the production of biological particles through growth that greatly increases local entropy production.

#### 2.1.3 Transport model

Siders Pond is horizontally well mixed, so an advection-dispersion-reaction (ADR) model that includes particle sinking was used to approximate vertical transport of the 19 state variables (Fig. 1). The origin of the vertical coordinate, *Z*, is defined at the pond’s surface, and the axis points in the positive direction downward towards the benthos and reaches a maximum depth of 15 m. Siders Pond 3D bathymetry surface (Fig. S1) was rendered from a contour plot in Caraco (1986), and an equation for cross-section area as a function of depth, *A*(*Z*), was derived therefrom (Eq. S3). Equations for vertical volumetric flow rate, *q*(*Z*), lateral groundwater inputs, *q_L_*(*Z*) (Eq. (S5)), and seawater intrusion at the bottom boundary were determined from Caraco (1986), who used both tritium-helium-3 dating combined with mass balance calculations to estimate freshwater inputs and seawater intrusion. An equation for the dispersion coefficient as a function of depth, *D*(*Z*), (Eq. (S6) and Fig. (S2)), was derived by fitting model predicted salinity profiles to observations collected during this study. Salinity profiles predicted by the ADR model show good agreement to observations (Fig. S2).

The primary external drivers in the model are temperature, *T*(*Z*), pH, *p_H_*(*Z*), and photosynthetic active radiation (PAR), *I*(*t, Z*). Since the ADR model does not include either energy nor proton balances, equations for both *T*(*Z*) and *p_H_*(*Z*) were simply based on linear interpolations of observations obtained from the water quality probe used during sampling. Surface irradiance, *I*(*t*,0), was based on a model of solar zenith angle (Brock, 1981), which assumes a cloudless sky. To predict PAR light intensity as function of time and depth, a standard light adsorption model was used that includes coefficients for light absorption by water, *k_w_*, and particulate material, *k_p_* (Eq. S7).

To complete the ADR transport model, Neumann boundary conditions were used for state variables at the pond’s surface, but particles were not allowed to sink into the boundary from above (Eq. S9), and gas transport for O_2_, CO_2_ and H_2_S across the air-water interface was accounted for using a stagnant film model (Eq. S10). Robin boundary conditions were used at the bottom boundary based on the flux of material entering the boundary associated with the intrusion of seawater diluted with groundwater from below. However, aerobic and anaerobic decomposition of sinking organic matter from the water column contributed to a sink for O_2_ and H_2_SO_4_, and a source for H_3_PO_4_, H_2_CO_3_ and H_2_S to the overlying water (Eq. S23).

##### 2.1.3.1 Parameter adjustments

One of the objectives in using a variational approach is to remove model parameters, but a few required adjustments to get simulations to within an order of magnitude of Siders Pond observations. Our objective was not to minimize error between predictions and observation, so the few parameters described in this section were adjusted in a rather ad hoc manner. No formal data assimilation was attempted, and the parameters were adjusted using fixed nominal values for the optimal control parameters, *ε_j_* and Ω_*i,j*_ (i.e., simulations without optimization). All biological structures, 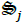, detritus, C_*D*_ and P_*D*_, and associated phytoplankton carbohydrates, C_*Phy*_ and C_*GSB*_, were allowed to sink through the water column (Eq. (S1)). However, sinking velocities can vary greatly across and within organism size classes (Bach et al., 2012), so velocities where set through trial and error to obtain profiles consistent with those observed in Siders Pond (Table S1). In addition, coefficients for light absorption by water, *k_w_*, and particles, *k_P_*, were adjusted so that light profiles were similar to observations (SM Section 2.5). Computational approach used for solving the ADR equation is described in *SM* Section 2.9.

#### 2.1.4 Optimization and MEP

The stoichiometry, thermodynamics and kinetics of the 28 reactions that comprise the metabolic network (Table 1, Fig. 1, SM Section 2.7) vary as a function of the *ε_j_* optimal control variables, and the partitioning of biological structure, 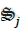, to sub-reactions *r_i,j_* (Table 1) that depends on the values of the Ω_*i,j*_ control variables. A solution to the MEP model, and associated microbial biogeochemistry, is determined by adjusting *ε_j_* and Ω_*i,j*_ over time and space to maximize entropy production, the details of which are provided in *SM* Section 3.2, but also see Vallino et al. (2014). The critical aspect of the optimization is choosing the appropriate time interval over which to maximize entropy production and whether local or global optimization should be used (Vallino, 2011); consequently, these two aspects are the focus of this manuscript and form the bases of the Results Section. Computational approach used for solving the optimal control problem is described in *SM* Section 3.2.

### 2.2 Siders Pond site description

Siders Pond is a small coastal meromictic kettle hole (volume: 10^6^ m^3^; area: 13.4 ha; maximum depth: 15 m) that receives approximately 1×10^6^ m^3^ of fresh and 0.15 x 10^6^ m^3^ of saltwater each year (Caraco, 1986). The latter input occurs via episodic inputs during high tides and storm events via a small creek that connects the pond to Vineyard Sound approximately 550 m to the south (Fig. S1). Tritium-helium water dating confirmed vertical mixing across two observed chemoclines, but permanent stratification is maintained because the saltwater inputs enter the pond at depth, mix upward and become entrained with freshwater before exiting the pond (Caraco, 1986). Caraco (1986) also characterized N and P loading to the pond (50 g N m^2^ y^-1^ and 1.3 g P m^-2^y^-1^, respectively), and an N+P enrichment study (Caraco et al., 1987) shows phytoplankton to be P limited, especially in the low salinity surface waters. Previous studies show Siders Pond is eutrophic averaging 16 mg m^-3^ chlorophyll a (Chl a) in surface waters (but can exceed 100 mg m^-3^ at times) and an annual primary productivity of 315 g C m^-2^ (Caraco, 1986; Caraco and Puccoon, 1986). In anoxic bottom waters bacterial Chl c, d and e associated with photosynthetic green sulfur bacteria averages 20 mg m^-3^ (purple sulfur bacteria were not found in high concentration), but BChl cde was also observed to reach high concentrations at times (> 75 mg m^-3^). Even though green sulfur bacteria could attain high concentrations, their productivity was only 6% of the oxygenic photoautotroph (cyanobacteria + algae) production (Caraco, 1986). Siders Pond was chosen for this study because extensive redox cycling occurs over a 15 m deep water column, which greatly facilitates sampling due to the large water volumes that can be readily collected without perturbing the system.

### 2.3 Sampling and measurements

Samples were collected from Siders Pond, Falmouth, MA over a 24 h period starting at 6:45 on Jun 25^th^ and ending at 7:37 on Jun 26^th^, 2015 from a single station located within the deepest basin of the pond (41.548212°N, 70.622412°W). A total of 7 casts were conducted over the 24 hr period, and each cast sampled 8 depths to generate a 2D sampling grid designed for comparison to model outputs (Fig. 2). A Hydrolab DS5 water quality sonde (OTT Hydromet, GmbH) was connected to a Hydrolab Surveyor 4 handheld display and used to record depth, temperature, salinity, pH, dissolved oxygen (DO), photosynthetic active radiation (PAR) and in situ Chl a fluorescence. All sensors were calibrated per manufacturer’s instructions one day prior to sampling. One end of a 20 m long section of vinyl tubing with a 1 cm inside diameter was attached to the water quality sonde, while the other end passed through a Geopump 2 (Geotech, CO) peristaltic pump and then connected to 25 mm polypropylene, acid washed in-line Swinnex filter holder (Millipore, MA), which housed a GF/F glass fiber filter (Whatman, GE Healthcare) that had been ashed at 450°C for 1 hr. At all depths, the vinyl tubing was first flushed for at least 2 min, sample collection vials were washed twice with filtrate and GF/F filters where changed as needed to maintain high flow. This design allowed water samples to be collected at the desired depths and processed on location then preserved on ice or dry ice for later analysis as described below.

**Fig. 2.**
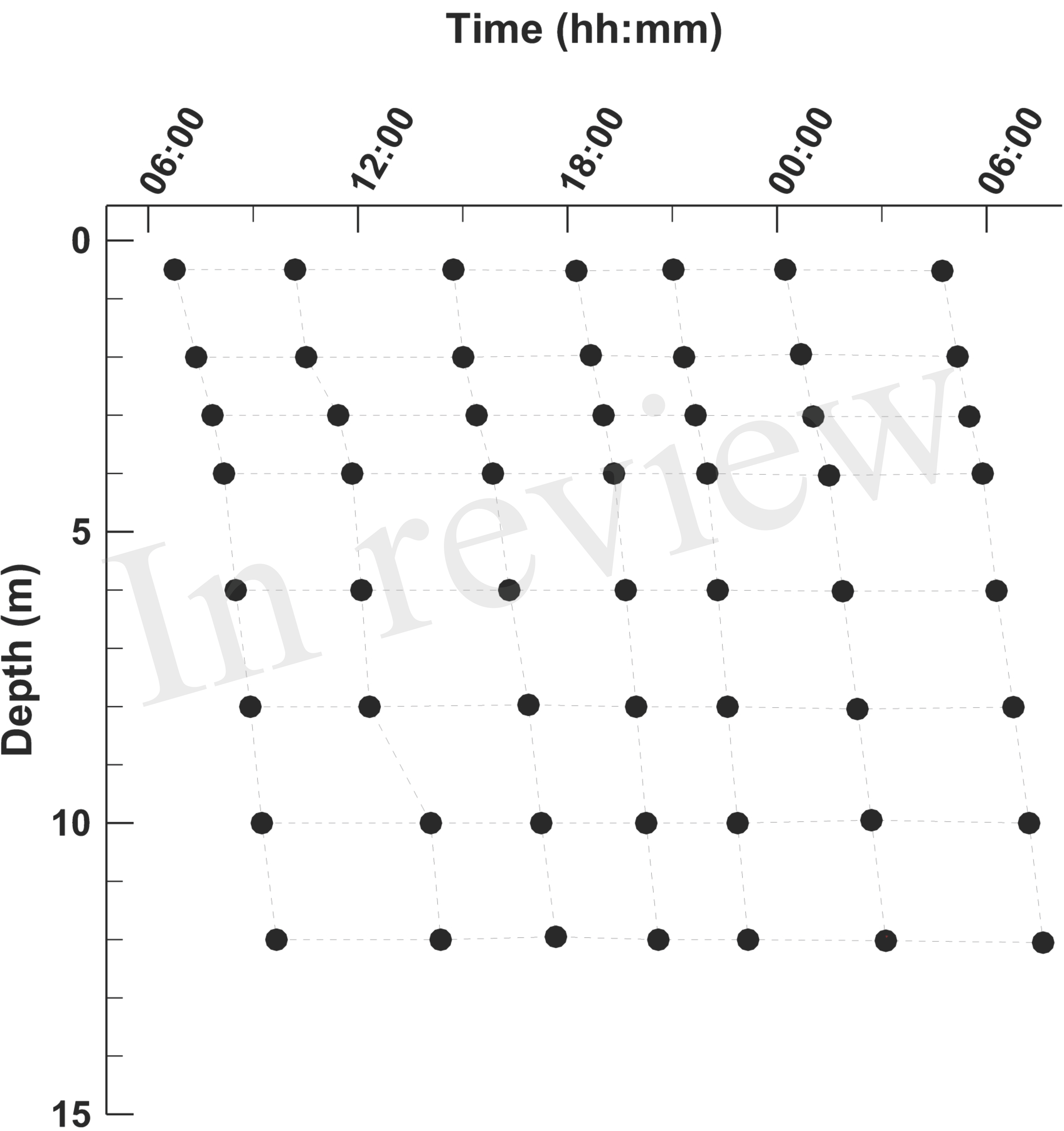
Layout of the Siders Pond 2D (*t,z*) sample grid. Samples were collected over a 24 h diel cycle starting at ~ 6:00 on 25 Jun 2015 and ending at ~6:00 on 26 Jun 2015. Samples were collected at 0.5, 2, 3, 4, 6, 8, 10 and 12 m.

Unless otherwise noted, all samples were filter as described above and stored in 20 mL high-density polyethylene scintillation vials (Fisher Scientific). Samples were preserved for later analysis as follows. Inorganic phosphate: 15 mL samples were amended with 20 μL of 5 N HCl and placed on dry ice. Dissolved inorganic carbon (DIC) and sulfate: samples were collected in 12 mL exetainers (Labco, UK) by overfilling bottles from the bottom up, and then capped without bubbles and placed on ice. Hydrogen sulfide: while filling exetainers, 25 μL of sample was pipette transfered to 6 mL of 2% zinc acetate in a 20 mL scintillation vial and place on ice. Dissolved organic carbon (DOC): 25 mL of sample was collected in previously ashed 30 mL glass vials to which 40 μL of 5 N HCl was added then stored on ice. Particulate organic carbon (POC): approximately 300 mL of sample was passed through new, ashed, 25 mm GF/F filter then stored in a plastic Petri dish on dry ice.

Samples were analyzed at the Ecosystems laboratory, MBL as follows. Phosphate: samples were stored at -20°C then later analyzed following the spectrophotometric method of Murphy and Riley (1962) on a UV-1800 spectrophotometer (Shimadzu, Kyoto, Japan). DIC: samples were immediately run on return to MBL on an Apollo AS-C3 DIC analyzer (Apollo SciTech, DE). Sulfate: samples were sparged with N_2_ to strip H_2_S on return to MBL and stored at 4°C then analyzed using ion chromatography on a Dionex DX-120 analyzer (Dionex, Sunnyvale, CA). Hydrogen sulfide: samples were briefly stored at 4°C for five days then analyzed using the spectrophotometric method of Gilboa-Garber (1971). DOC: samples were stored at 4°C then run on a Shimadzu TOC-L high temperature total organic carbon analyzer at 720°C. POC: samples were stored at -20°C then analyzed on a Thermo Scientific FLASH 2000 CHN analyzer using aspartic acid standards. To facilitate model comparison to observations, Chl a was estimated from modeled output of phototroph biomass concentrations using 12. 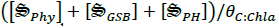, where 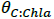 is the C:Chl a ratio, which was set to 50 μg C (μg Chl a)^-1^ (Sathyendranath et al., 2009).

## 3 Results

During initial model develop, a simple study was conducted to compare local versus global MEP optimization (Vallino, 2011) for the Siders Pond model; however, it was quickly realized that global optimization did not produce relevant solutions (see Discussion Section for explanation). Consequently, all solutions presented here are from local entropy maximization at 10 depths (0, 1, 2, 3, 4, 6, 8, 10, 12, and 15 m) conducted in parallel. We investigated two optimization time intervals; ∆*t*, for entropy maximization, over a short interval of 0.25 days as well as over a long interval of 3 days. To minimize the influence of uncertain initial conditions on final solutions, both the short and long interval optimizations were started on May 19^th^ and ran with short or long interval optimization until Jun 21^st^. Solutions from Jun 21^st^ were then used as initial conditions for 6 day production simulations that required 24 intervals for the short 0.25 d optimization and 2 intervals for the long 3 d optimization. The 6 day simulations ended on Jun 27^th^ to allow comparison with observations collected on Jun 25^th^ and 26^th^ (Fig. 2). Our results focus on how these two solutions compare to observations as well as to each other. Future discounting was not used in solutions presented here (i.e., *k_D_* = 0 and δ*t* = ∆*t*). See *SM* Section 3.2 for details on optimization.

### 3.1 Simulations compared to observations

In this section, solutions obtained from the short (0.25 d) and long (3 d) interval optimizations are compared to biogeochemical observations from Siders Pond that were collected on Jun 25^th^ and 26^th^, 2015 (Fig. 2). Simulated profiles for photosynthetic active radiation (PAR), Chl a and dissolved oxygen over 6 days for the short and long interval optimizations are compared to 1 day of observations in Siders Pond in Figure 3. The short interval optimization (SIO) shows PAR extending to nearly the bottom of the pond (Fig. 3a), while PAR from the long interval optimization (LIO) (Fig. 3b) more closely matches observations (Fig. 3c); however, high intensity PAR (~ 100 μmol m^-2^s^-1^) observations extend a few meters deeper than the LIO simulation predicts. The prediction for Chl a in both simulations do not match observations very well (Fig. 3d-f), but this is partially due to how Chl a was estimated in the model, since Chl a is not specifically modeled (see Section 2.3). The Chl a *in vivo* observations show a peak Chl a around 5 m, while both simulations have peaks around 12 m, but those peaks are due to accumulation of sinking phytoplankton rather than productivity (see below). The LIO simulation does show a secondary Chl a peak developing around 3 m, but it is weaker than observations (~ 6 vs 40 μg L^-1^). Based on DO, the SIO shows anaerobic conditions begin at 6 m, while the LIO shows that occurring at 10 m. Observations for the transition to anoxia splits between the two simulations at 8 m. Observations also show a subsurface DO maximum at 3.5 m, while both simulations show max DO closer to 1.3 m. Furthermore, the SIO shows a decreasing DO max with time, while the LTO shows an increase over time, and the maximum reaches 800 mmol m^-3^ versus 480 for observations. In general, the SIO simulations show less phototrophic activity while LIO shows greater activity than what the observations indicate.

**Fig. 3.**
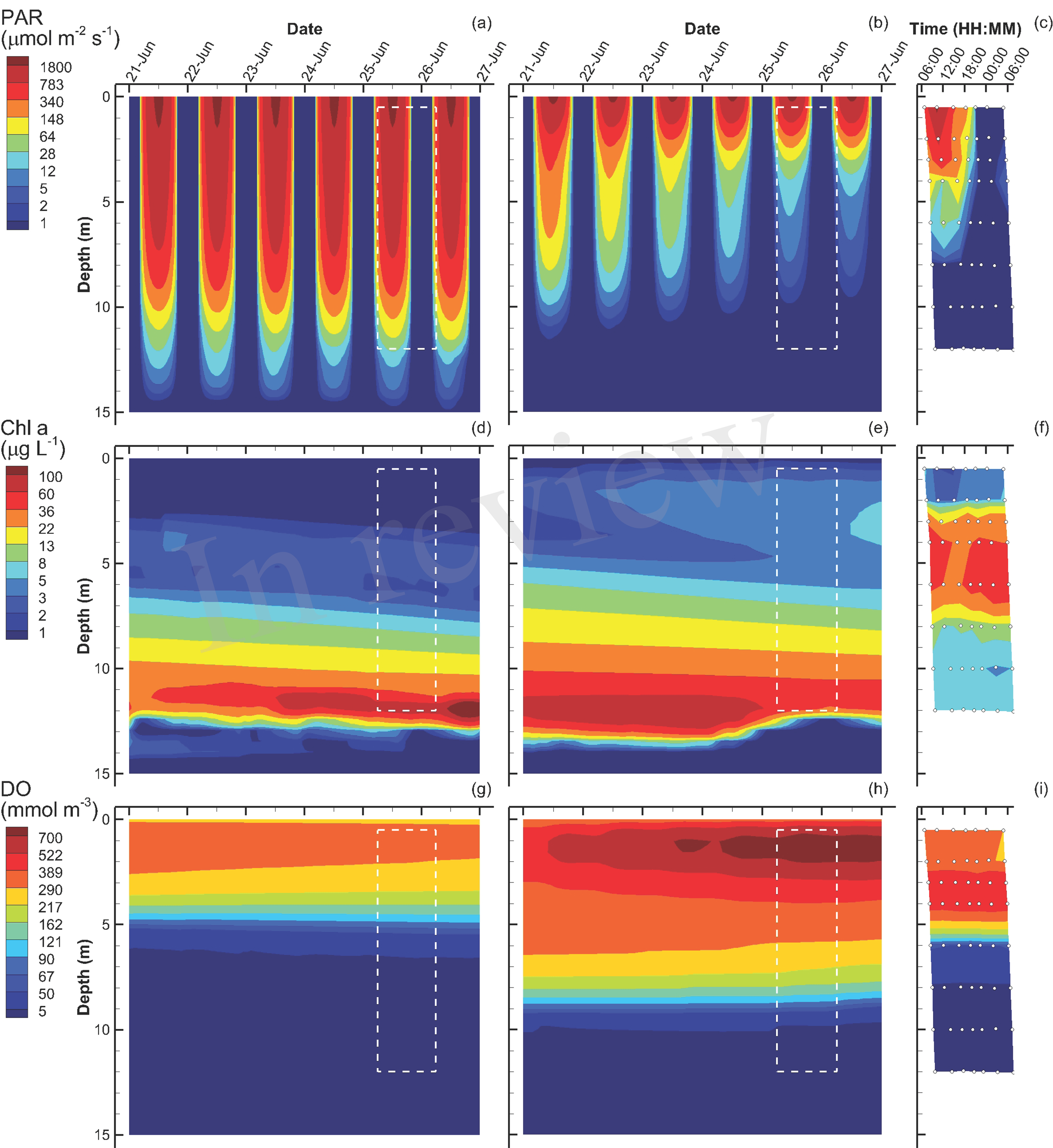
Contour plots of photosynthetic active radiation (PAR) (a-b), chlorophyll (Chl) a concentration (d-f) and dissolved oxygen (DO) (g-i) for six day simulations using short interval optimization (SIO) (left column), long interval optimization (LIO) (center column) and for observations collected from Siders Pond over a 24 hour period on Jun 25 to Jun 26 2015 (right column). Rectangle (dashed white lines) in simulation plots corresponds to time and depths where observations are comparable (i.e., Jun 25/26, 0.5 to 12 m). Actual observations are shown as white circles with black parameters (see also Fig. 2).

Simulations of substrate concentrations for autotrophs, namely inorganic phosphate and dissolved inorganic carbon (DIC), show overall agreement to observations for both SIO and LIO solutions (Fig. 4), but some discrepancies are apparent. The phosphate chemoclines occur around 10 m, 11 m and 8 m for the SIO, LIO and observations, respectively (Fig. 4a-c). Above the phosphate chemocline, the SIO simulation shows slightly elevated levels of H_3_PO_4_ (~0.3 mmol m^-3^) in the surface and 5 m layers, compare to observations (Fig. 4c), which are near the level of detection (0.03 mmol m^-3^) except for a few spikes. Below the phosphate chemocline, observations show slightly higher accumulations of phosphate, approaching 22 mmol m^-3^, while the SIO and LIO simulations show max concentrations closer to 16 and 19 mmol m^-3^, respectively. For DIC, both SIO and LIO simulations show much lower DIC concentrations below 10 m (3,000 and 5,400 mmol m^-3^, respectively) than observed in Siders Pond (16,500 mmol m^-3^), which would indicate either much higher anaerobic respiration in the pond than occurs in the simulations, or the bottom boundary condition for the simulations is incorrect (see *SM* Section 2.6). The SIO and LIO also show greater draw down of DIC in the surface water above 2 and 4 m respectively, and the LIO simulation shows minimum DIC approaching just 2 mmol m^-3^ at 1 m, while minimum observed value is above 660 mmol m^-3^.

**Fig. 4.**
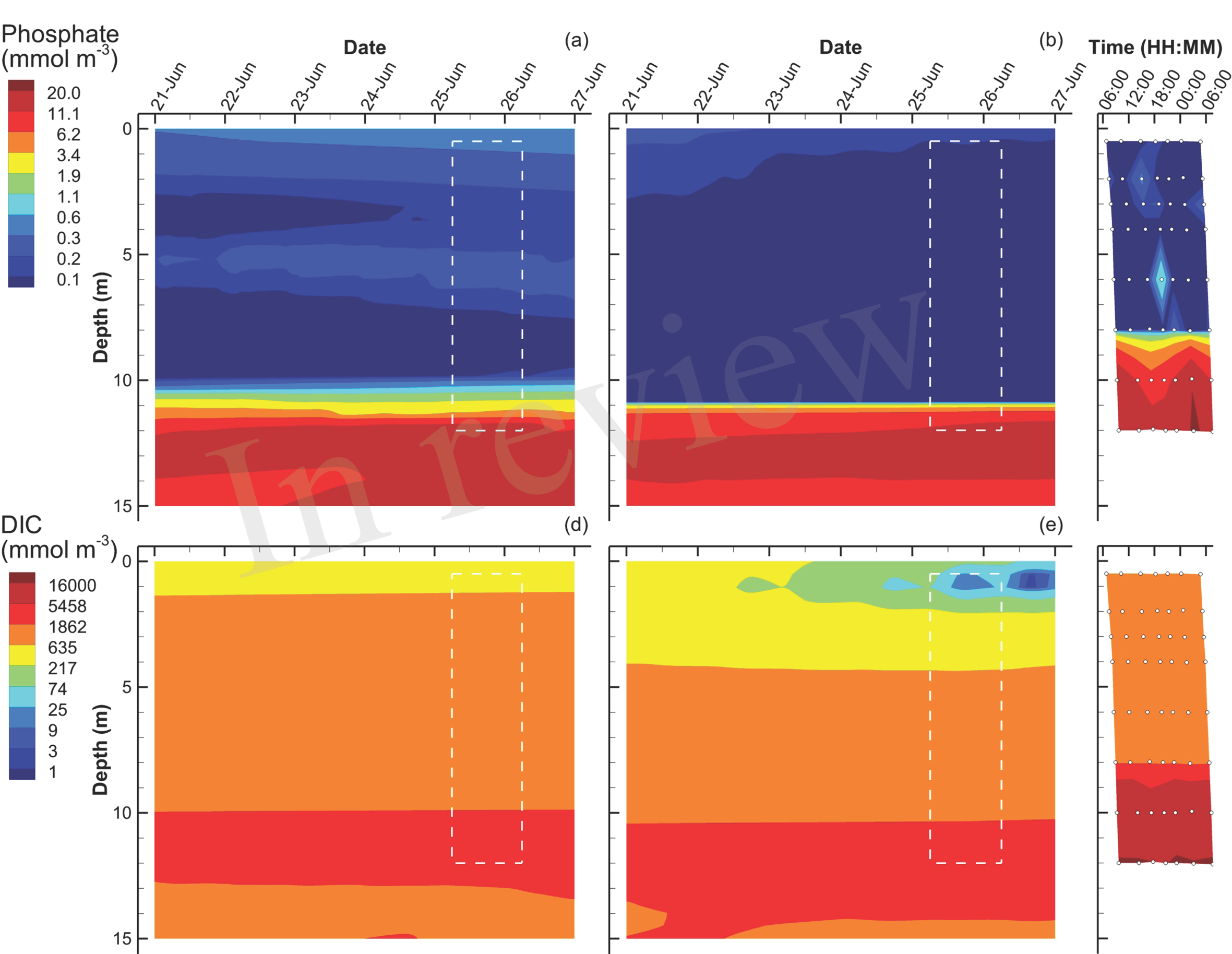
Contour plots of inorganic phosphate (a-c) and dissolved inorganic carbon (DIC) (d-f) concentrations for SIO and LIO simulations (left and center columns, respectively) and Siders Pond observations (right column). See caption to Fig. 3 for other details.

Simulations of hydrogen sulfide show a chemocline at approximately 9 and 10 m for the SIO and LIO solutions, respectively, which are comparable to the H_2_S chemocline observed in Siders Pond at about 8 m (Fig. 5a-c). However, H_2_S reaches concentrations as high as 7,000 mmol m^-3^ in Siders Pond, while maximum concentrations only reaches 900 and 2,100 mmol m^-3^ in the SIO and LIO simulations, respectively. Simulations also show a peak H_2_S concentration at 12 m, and a decrease in concentration below 12 m, which seems unlikely and indicates that the bottom boundary condition may not be correct. That is, the flux of H_2_S into the bottom water is lower than it should be, but it is also possible that the rate of anaerobic metabolism in the simulations is too low, which would be consistent with the low simulated DIC concentration compare to observations mentioned above. Simulated sulfate concentrations in the upper portion of the water column (0 to 4 m) are similar to those observed (Fig. 5d-e), but the simulations show a very dramatic sulfate chemocline starting at about 8 m, while observations show a more gradual increase in sulfate with depth. Furthermore, sulfate reaches much higher concentrations in the simulations at depth than do observations, showing maximums of 15,500 mmol m^-3^ in both simulations, while observation maximum is only 9,000 mmol m^-3^. The lower simulated concentrations of H_2_S and the higher simulated sulfate concentrations indicate that sulfate reduction in the model is lower than that actually occurring in Siders Pond.

**Fig. 5.**
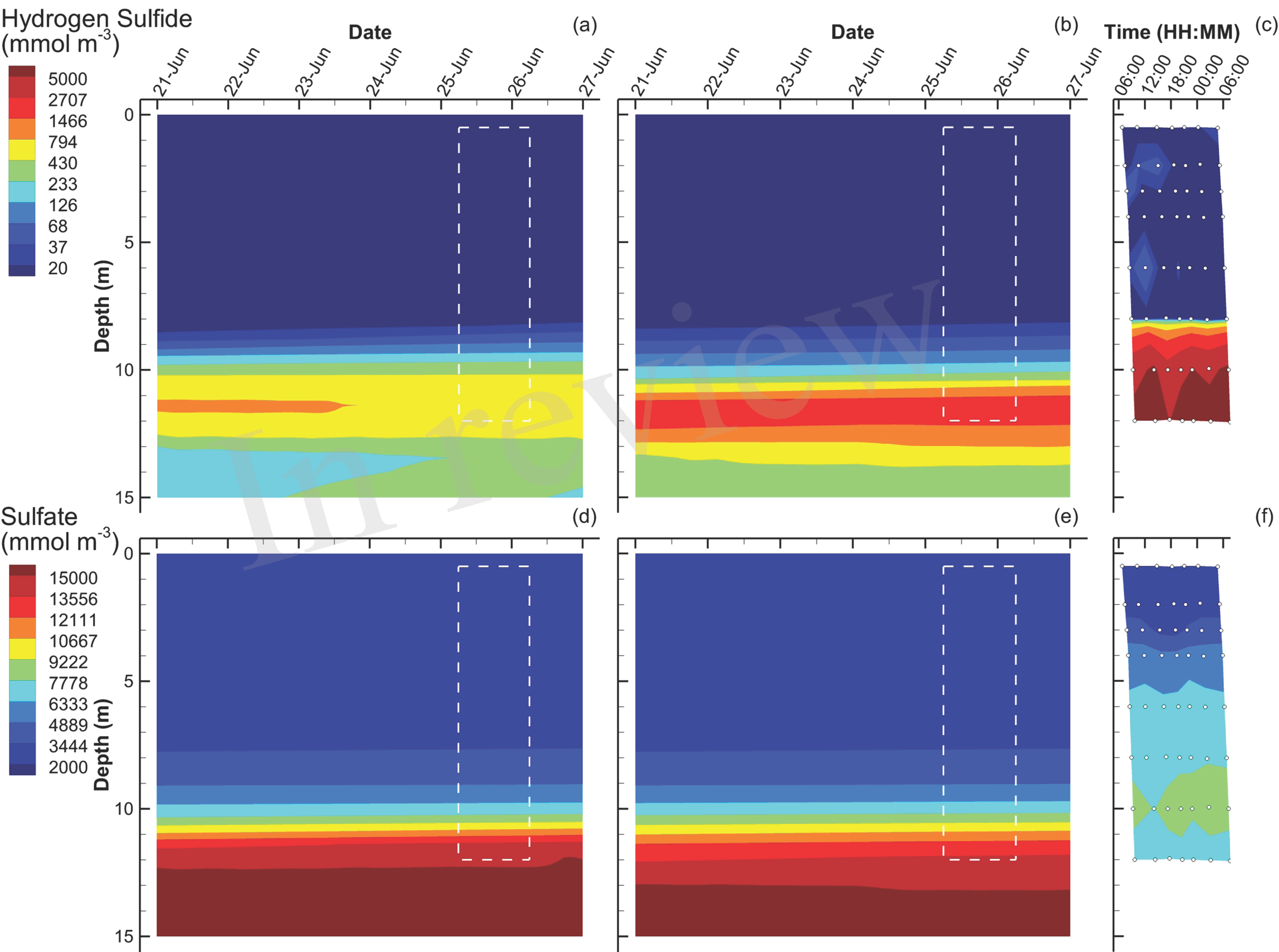
Contour plots of hydrogen sulfide (a-c) and sulfate (d-f) concentrations for SIO and LIO simulations (left and center columns, respectively) and Siders Pond observations (right column). See caption to Fig. 3 for other details.

The last of the observations are dissolved organic carbon (DOC) and particulate organic carbon (POC) concentrations (Fig. 6). For simulations, DOC is a derived quantity based on the sum of state variables [C_*L*_]and [C_*D*_], while POC is derived from the sum of [C_*Phy*_], [C_*GSB*_]and the concentrations of all biological structures, 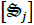. In the water column above 12 m, the SIO simulation shows very low concentrations of DOC (~ 1 mmol m^-3^), but then increases rapidly to a maximum of 1,900 mmol m^-3^ (Fig. 6a). The LIO simulation shows a similarly high DOC concentration at 14 m, but above 12 m, the DOC concentration ranges from 2 to 140 mmol m^-3^, which is comparable to those observed in Siders Pond, which range from 200 to 300 mmol m^-3^ above 10 m, and increase to a maximum of about 1,000 mmol m^-3^ at 12 m. Similar to DOC, POC in the SIO simulation shows low values (< 30 mmol m^-3^) above 6 m, but POC peaks to 1,000 mmol m^-3^ at 12 m (Fig. 6d). The POC concentrations from the LIO simulation are closer to observations, but the mid-water POC maximum in the LIO simulation is approximately 500 mmol m^-3^, while the observations peak at 280 mmol m^-3^ around 4 m. Like the SIO simulation, the LIO simulation also shows high POC concentrations below 10 m, which was not observed in Siders Pond. It is possible that DOC and POC may reach higher concentrations in the funnel-like basin of Siders Pond below 12 m (Fig. S1), but samples were not collected there.

**Fig. 6.**
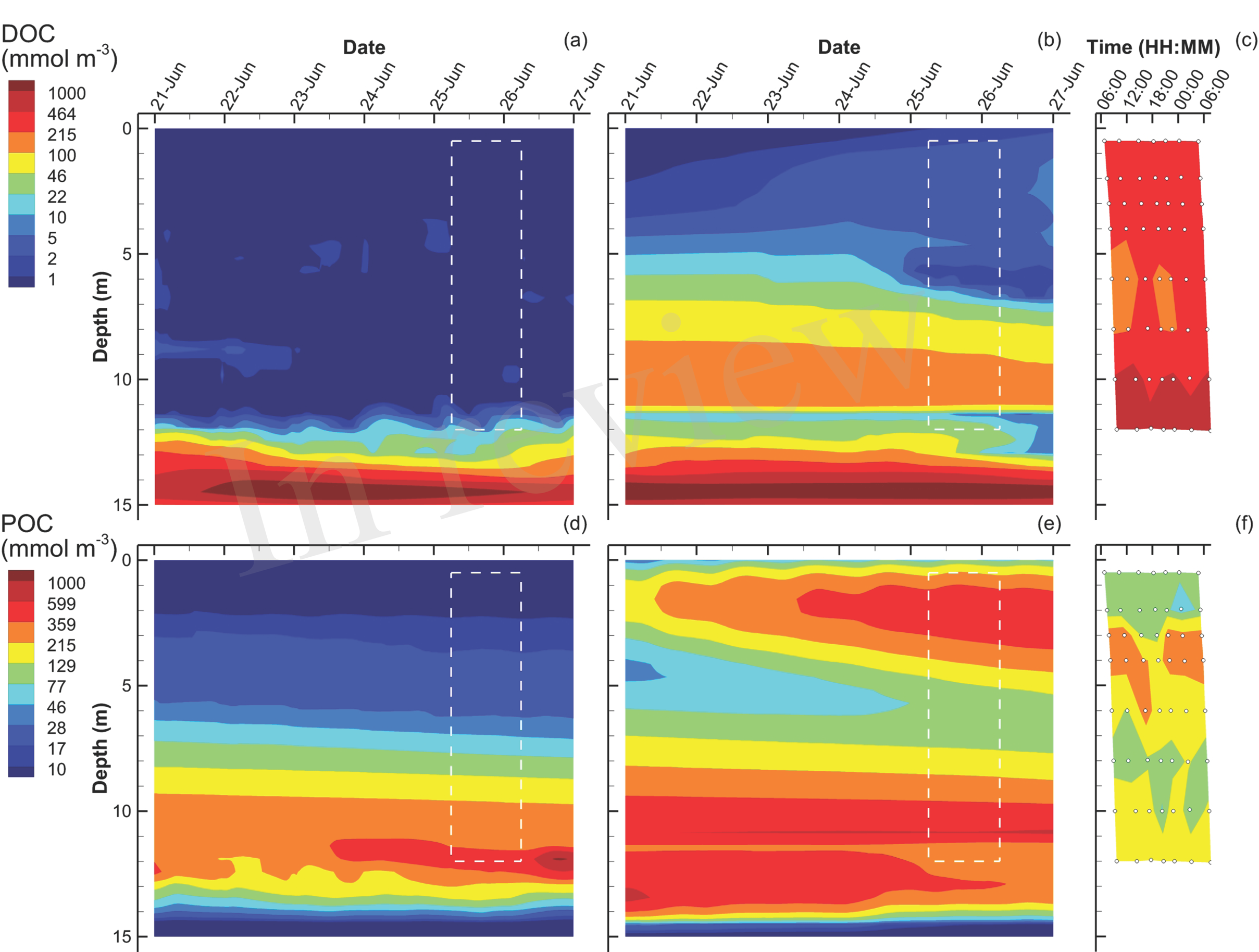
Contour plots of dissolved organic carbon (DOC) (a-c) and particulate organic carbon (POC) (d-f) concentrations for SIO and LIO simulations (left and center columns, respectively) and Siders Pond observations (right column). See caption to Fig. 3 for other details.

### 3.2 Comparison between SIO and LIO simulations

Since the MEP optimization model generates a large number of outputs, this section highlights some of those outputs to contrast the simulations based on the short (0.25 d) interval optimization (SIO) to that from the long (3 d) interval optimization (LIO). Consider entropy production (Eq. S178), which is the variable that is being sequentially maximized over either a 0.25 d interval (SIO) or a 3 d interval (LIO) at 10 different depths (see SM Section 3.1, Eq. (S180)) (Fig. 7). Total entropy production, 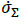, for the SIO and LIO simulations differ significantly, in that 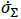 in the SIO solution is spread out over a 12 m water column (Fig. 7a) versus the LIO solution (Fig. 7e) where most of the entropy production occurs in the top 3 m of the pond. Furthermore, peak entropy production in the LIO simulation is 7.5 times great than the SIO solution (9.8 vs 1.3 GJ m^-1^ K^-1^ d^-1^), and the total integrated entropy produced over the water column and 6 d simulation, *σ_T_* (Eq. (S182)), was 21.3 GJK^-1^ and 36.5 GJ K^-1^ for the SIO and LIO simulations, respectively. For comparison, if the pond was sterile, *σ_T_* would equal 12.4 GJ K^-1^ from light absorption in the water column. The higher total integrated entropy, *σ_T_*, produced by the LIO simulation illustrates that extending the time scale results in greater entropy production, as has been previously shown in experimental mesocosms (Vallino et al., 2014).

**Fig. 7.**
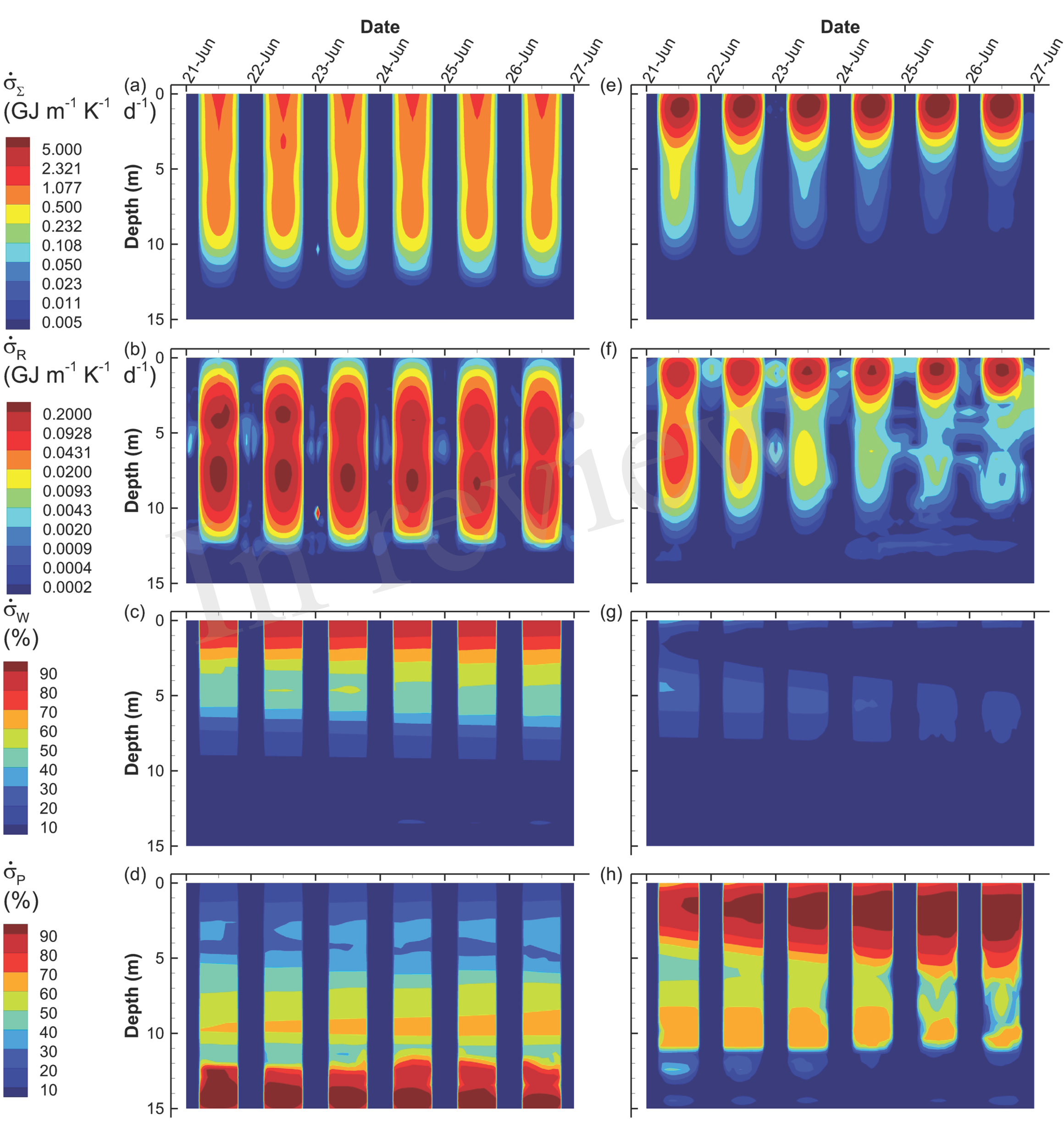
Contour plots of simulated total entropy production, 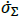 (Eq. (S178)) (a, e), entropy production from reactions, 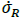 (Eq. (S178)) (b, f), and entropy production from light absorption by water, 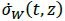 (Eq. (S176)) (c, g) and particles, 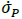 (Eq. (S177)) (d, h) for the SIO (left column) and LIO (right column) simulations. Entropy production from light dissipation by water and particles are shown as a percentage of total entropy production.

The different contributors to total entropy production, namely by reaction, 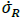, water, 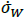, and particles, 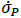, also differ significantly between the two simulations. For instance, total integrated entropy production associated with reactions (Table 1) was actually greater in SIO than the LIO simulation (Fig. 7b vs f), as well as exhibited a greater maximum 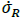 (0.28 vs 0.26 GJ m^-1^K^-1^d^-1^ for SIO versus LIO); however, entropy production by reaction during the day is rather small (< 25% in the upper 5 m) compared to light dissipation by water or particles, but the two simulations differ here as well. The SIO simulation dissipates most of the incoming radiation by water absorption in the upper 5 m of the water column (Fig. 7c vs g), while the LIO simulation dissipates most of the electromagnetic potential via absorption by particles (Fig. 7d vs h). As evident in the POC (Fig. 6) and Chl a (Fig. 3) concentrations, the LIO simulation produces more biomass in the upper portion of the water column, and biomass is effective at absorbing and dissipating light. The high biomass in the LIO simulation is due to phytoplankton, as describe next.

An analysis of phytoplankton growth by the SIO and LIO simulations (Fig. 8) illustrates not only how the two simulations differ, but also some of the mechanics of the MEP-based optimization approach. Phytoplankton density attains a maximum of 20 mmol m^-3^ in the LIO simulation by June 27^th^, but Phy are effectively absent in the SIO simulation, attaining a maximum of only 0.1 mmol m^-3^ (Fig. 8a and f). While it is possible that the low phytoplankton (Phy) density in the SIO simulation could be due to extensive predation, this is not the case because the rates of CO2 fixation (*r_1,Phy_*, Table 1) and conversion of fixed C to biomass (*r_2,Phy_*, Table 1) are two orders of magnitude lower in the SIO versus LIO simulation (Fig. 8b, c versus g, h). Of course the differences in phytoplankton density and reaction rates between the SIO and LIO simulations are due to how the optimal control variables change over time and space (Fig. 8d, e, i and j). Consider how reaction efficiency, *ε_Phy_*(*t,Z*), varies over time and space in the two simulations (Fig. 8d and i). The SIO simulation shows rapid switching between very high efficiencies (>0.98) and very low efficiencies (<0.02) over time. While not on all days, reaction efficiency drops to very low levels around noon (Fig. 8d), which results in high entropy production, but at the sacrifice of fixing CO_2_, which is consistent with a short term optimization objective: burn fast now instead of investing for the future. On the contrary, the LIO solution (Fig. 8i) shows much more gradual changes in *ε_Phy_*(*t,Z*), operating between 0.3 and 0.4 for most of the simulation. There is rapid changing of ε_*1,Phy*_(*t,Z*) in the LIO solution (Fig. 8j), but this makes sense, because the control variable partitions phytoplankton biomass to the light requiring carbon fixation reaction, *r_1,Phy_*, during peak daylight, then switches to the biosynthesis reaction, *r_2,Phy_*, at night (Fig. 8h). Changes in biomass partitioning is also observed in the SIO solution (Fig. 8e), but sometimes ε_*1,Phy*_ selects for the light-requiring CO_2_ fixation reaction at night. This nonsensical behavior is likely due to the nature of optimization. For the SIO simulation, phytoplankton do not contribute significantly to free energy dissipation, so values of the optimal control variable are freer to randomly fluctuate because they do not strongly impact the optimization objective. Based on Fig. 7c, the SIO solution places more weight on using water for the short term dissipation of electromagnetic potential in the pond’s surface, but in deeper water the SIO solution does grow biomass.

**Fig. 8.**
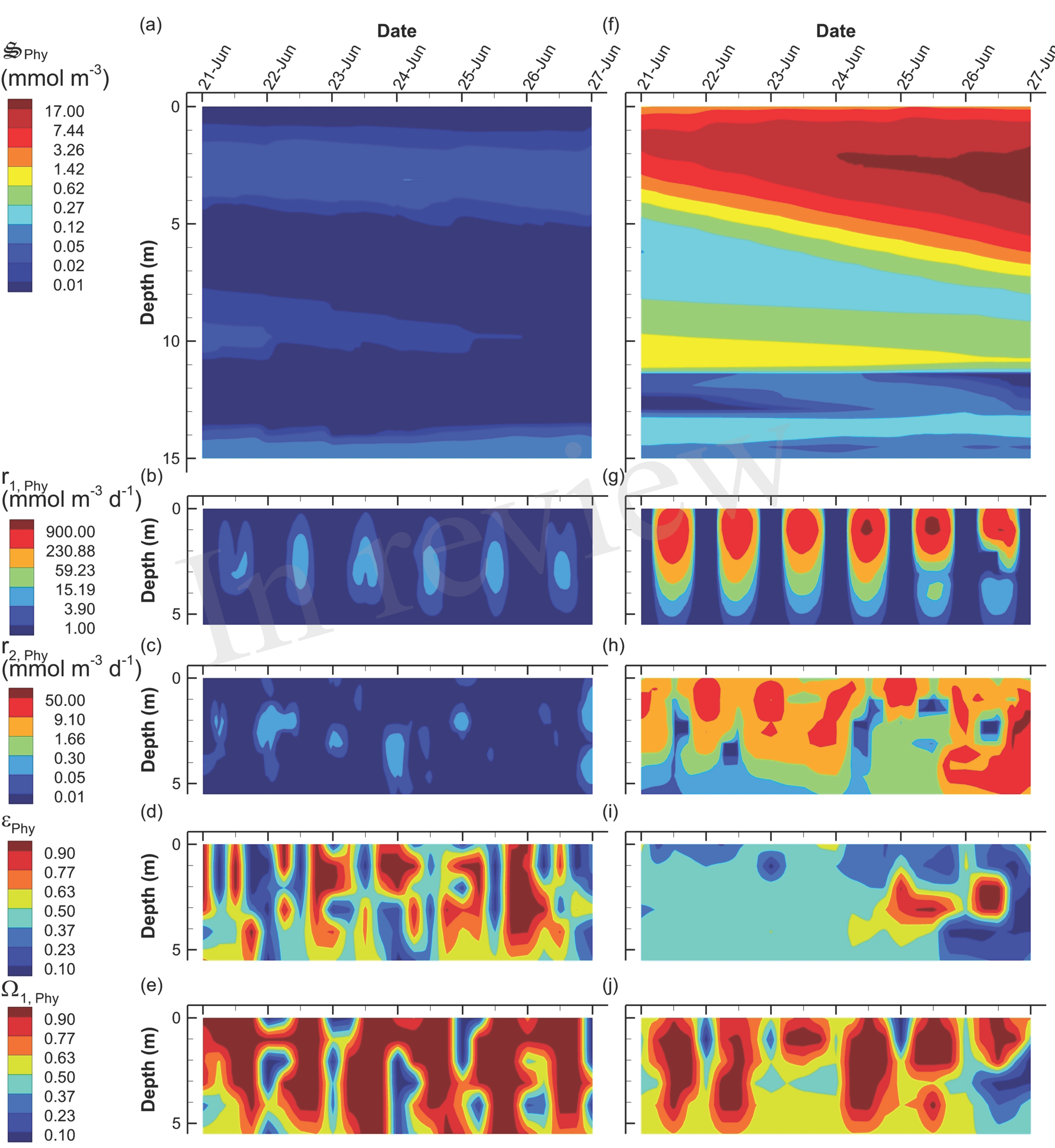
Contour plots of phytoplankton concentration, 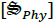 (a, f), rates for the two associated reactions, *r_1,Phy_* (b, g) and, *r_2,Phy_* (c, h) and two optimal control variables, *ε_Phy_* (d, i), and Ω_*1,Phy*_ (e, j) for the SIO (a-e) and LIO (f-j) simulations. Note, reaction rates and control variables are only plotted for the top 5 m of the water column where the processes are significant.

Instead of growing phytoplankton, the SIO solution produces more green sulfur bacteria (GSB), which reach a maximum concentration of 735 mmol m^-3^, compare to only 23 mmol m^-3^ in the LIO solution (Fig. 9a and d). Furthermore, GSB increase during the SIO simulation, while they decrease in the LIO, which is evident in the greater biosynthesis reaction, *r_2,GSB_* in SIO versus LIO solutions (Fig. 9b and e) as well as in *r_1,GSB_* (not shown). However, the value of the reaction efficiency control variable, *ε_GSB_*(*t,z*), does switch to low values around noon on several days in the SIO simulation, which implies again the solution favors entropy production over growth (Fig. 9c and f). This SIO simulation also favors much higher reaction rates of photoheterotrophs (PH), *r_1,PH_*, compared to the LIO solution, but interestingly, this does not lead to greater concentrations of 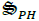 (Fig. 10). The reason is because high rates for *r_1,PH_* are coupled with extremely low values of *ε_PH_* (Fig. 10b and c). Based on the adaptive Monod equation that changes substrate affinity as a function of 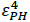 (Eq. (S131)), uptake of labile organic carbon by PH can occur at extremely low concentrations of C_*L*_ when *ε_PH_* is close to zero, but small *ε_PH_* values means very little biomass is produced as a results (Eq. (S123)). This is an interesting result, as light energy is being used to scavenge organic carbon at low concentrations, which contributes to the low labile organic carbon, C_*L*_, observed in the SIO simulation (Fig. 6a). Furthermore, the high entropy production from reaction, 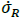 (Fig. 7b), is almost entirely due to PH. One of the reasons both green sulfur bacteria and photoheterotrophs are more active in the SIO versus LIO simulations is that light is not being intercepted by phytoplankton like it is in the LIO simulation (Fig. 8a vs f).

**Fig. 9.**
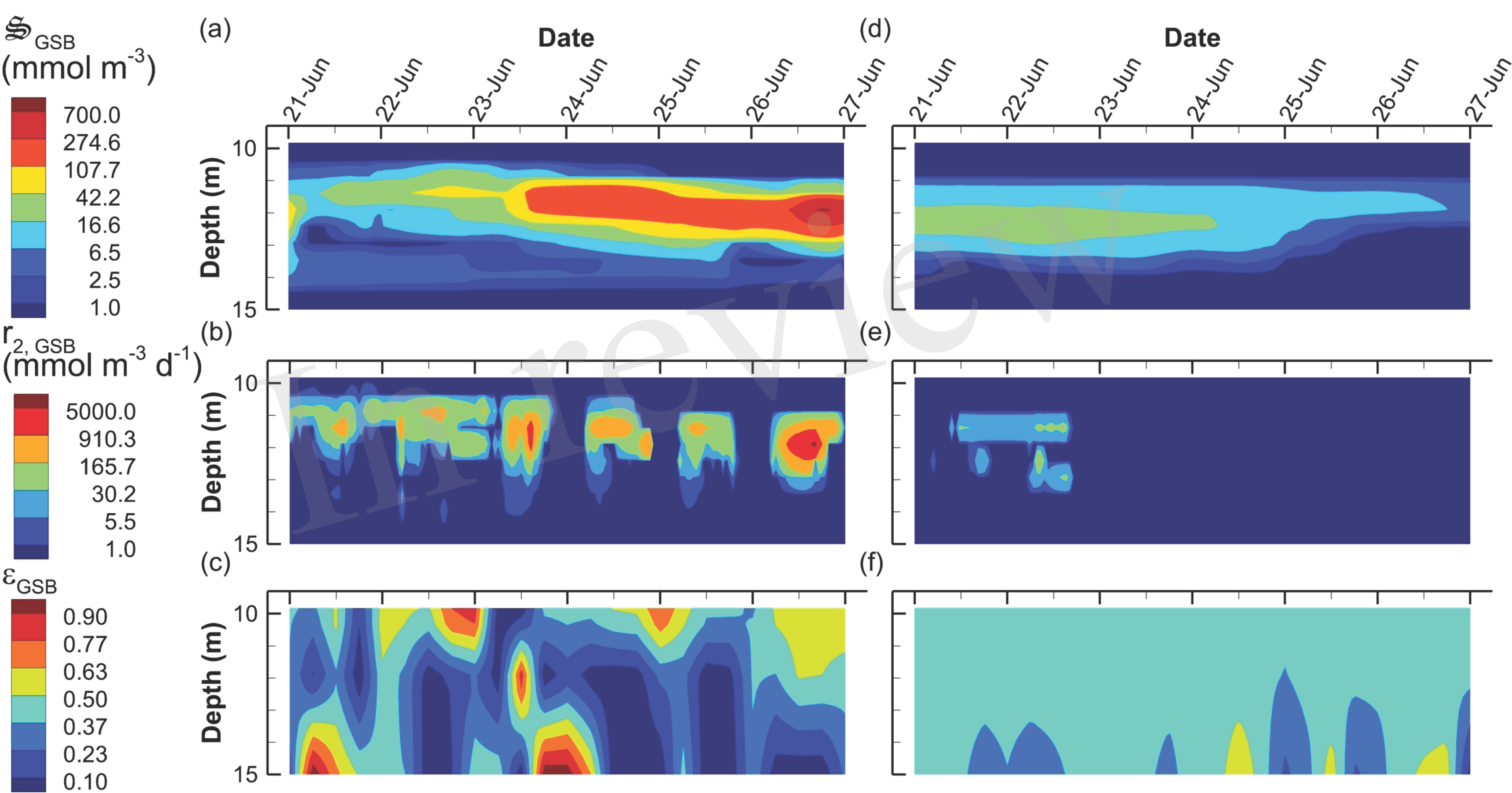
Contour plots of green sulfur bacteria concentration, 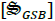 (a, d), rate of the biosynthesis reaction, *r_2,GSB_* (b, e) and the growth efficiency optimal control variable, *ε_GSB_* (c, f) for the SIO (a-c) and LIO (d-f) simulations. Note, variables are only plotted for 10 m to 15 m where the GSB are active.

**Fig. 10.**
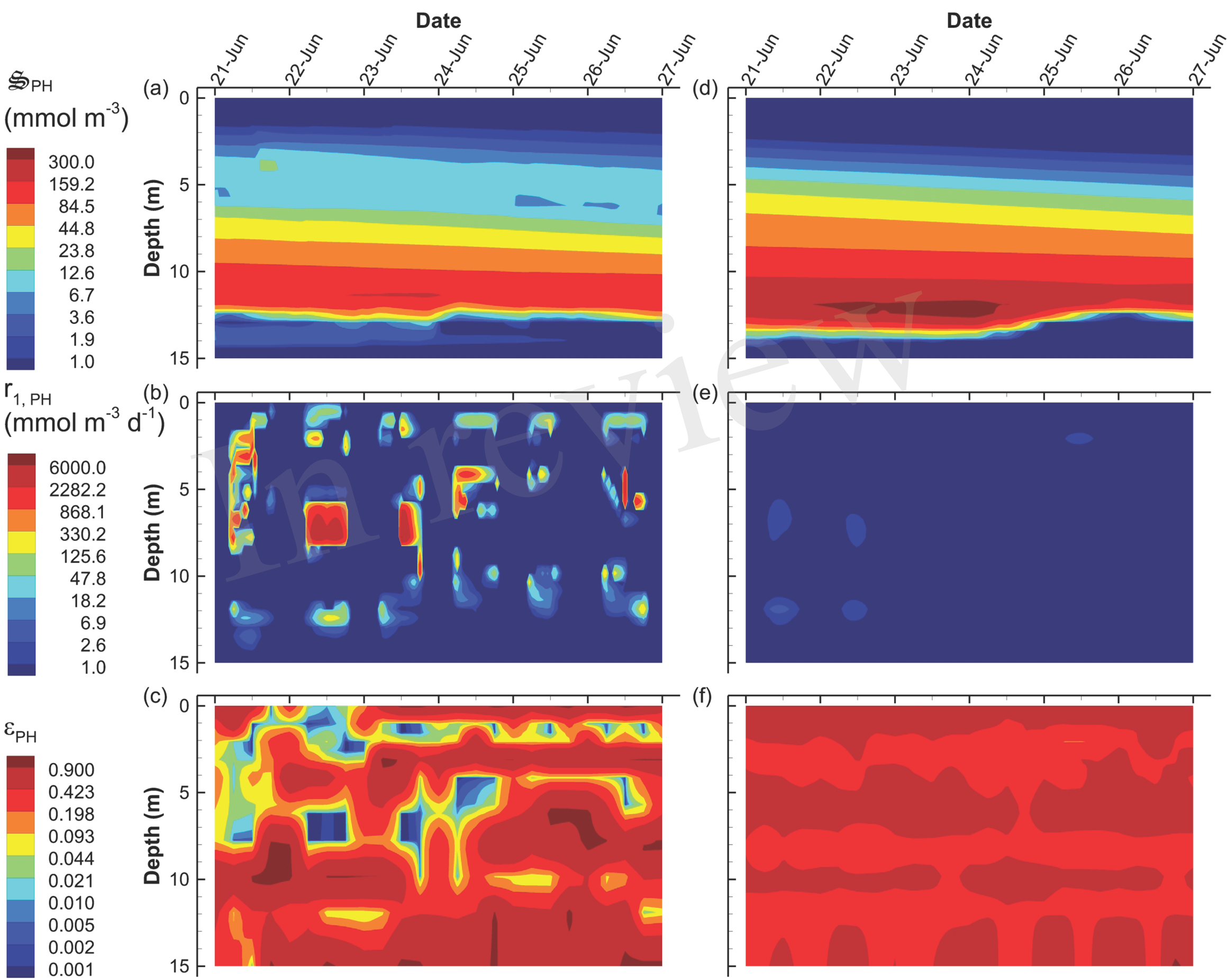
Contour plots of photoheterotroph concentration, 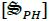 (a, d), rate of the carbon fixation reaction, *r_1,PH_* (b, e) and the growth efficiency optimal control variable, *ε_PH_* (c, f) for the SIO (a-c) and LIO (d-f) simulations. Note, contours for *ε_PH_* were selected to emphasize *ε_PH_* values near zero.

Bacterial densities are similar in the two simulations (Fig. S4a and d), but there is significantly higher growth rate by bacteria in the LIO simulation between 4 and 10 m (Fig. S4b and e). Both simulations allocate almost all of 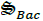 to detrital carbon decomposition, *r_2,Bac_*, below 13 m (Fig. S5c and f), but the SIO simulation also allocates biomass to *r_2,Bac_* sporadically throughout the water column to produce C_*L*_ (Fig. S4c), which is limiting (Fig. 6a). The anaerobic, sulfate reducing bacteria function similarly as bacteria, but of course operate in the anaerobic portion of the water column (Figs. S6 and S7). However, while their growth occurs predominately below 12 m (Fig. S6b and e), there is significant biomass accumulation at and above 11 m, which indicates advection is transporting them upwards. Like bacteria (Bac), the sulfate reducing bacteria (SRB) allocate biomass to *r_2,Bac_* below 13 m in both simulations (Fig. S7c and f) and sporadically throughout the water column in the SIO simulation (Fig. S7c). Overall, bacteria and SRB function in a complementary mode across the aerobic and anaerobic portions of the water column. Similar to phytoplankton, the control variables for both 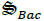 and 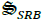 show more rapid (bang-bang) control in the SIO compared to the LIO simulation (Figs. S5 and S7). The sulfide oxidizing bacteria, 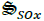, are largely unimportant in either of the 6 day simulations.

Another significant difference between the SIO and LIO simulations is a greater importance of predation, 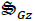, in the SIO solution (Fig. 11). Because predation is abstracted in the MEP model, it represents all predation mechanisms, including protists, predatory bacteria, viruses and cannibalism. In addition to dissipating chemical potential stored as biomass, predation serves a more important task of turning over biomass locked into metabolic functions, such as sulfate reduction, that are no longer of use under prevailing conditions. The SIO and LIO simulations show that the concentration of 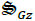 is more the 4 times higher in SIO than LIO solutions, and 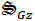 increases over time with the SIO objective. The primary prey items in the SIO solution are 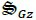 (i.e., cannibalism), 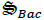 and 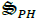 (Fig. 11b, c and d), while only 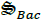 predation is of significance in the LIO solution (Fig. 11g).

**Fig. 11.**
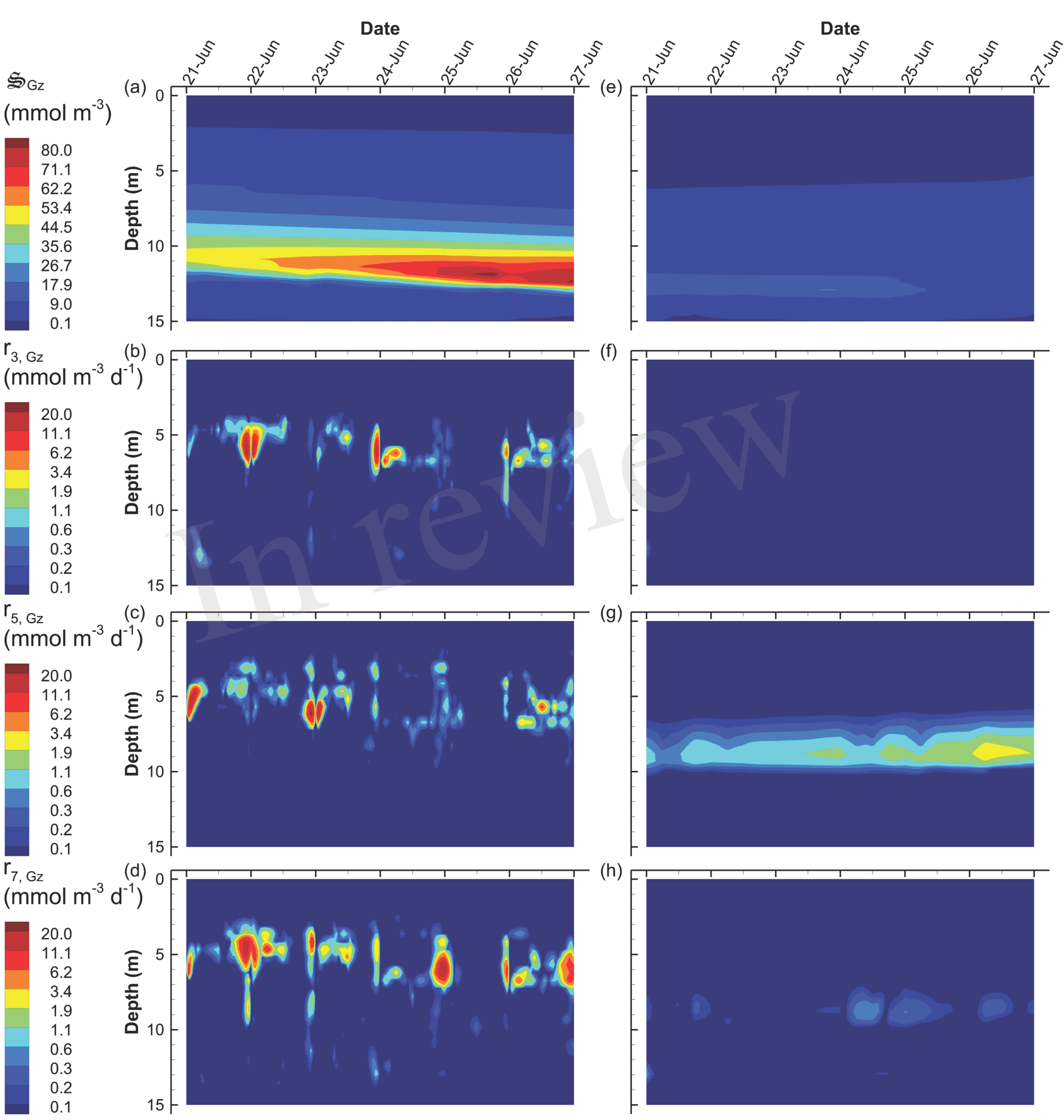
Contour plots of grazer concentration, 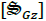 (a, e) and rates of predation on, grazers, *r_3,Gz_* (b, f), bacteria, *r_5,Gz_* (c, g), and photoheterotrophs, *r_7,Gz_* (d, h) for the SIO (a-d) and LIO (d-h) simulations.

Closer inspection of Fig. 11b-c reveals that predation occurs predominately at night, which is a result of temporal changes in the partitioning control variables, ε_*i,Gz*_, rather than prey concentration (not shown). It is also worth noting that while the high concentration of 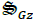 is found below 10 m, predation actually occurs between 4 and 7 m, which indicates the high 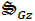 concentration is due to sinking and accumulation. Accumulation of biomass in the deep, anaerobic, portion of the water column becomes food for anaerobic predators, 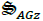, which are important in both SIO and LIO simulations, but are slightly more active in the LIO simulation (Fig. S8). The dominate prey items for AGz are GSB (Fig. S8b and e), AGz (cannibalism) (Fig. S8c and f), SRB (Fig. S9a and c) and PH (Fig. S9b and d).

An interesting result that derives from the focus on energy dissipation rather than organisms is the importance of chemotrophs on dissipating electromagnetic potential. For instance, heterotrophic bacteria, 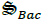, are well understood as dissipaters of chemical potential as they typically respire more reduced organic carbon than converting it biomass (i.e., *ε_Bac_* < 0.5, Eq. (S84)), where the latter reaction conserves chemical potential (Table 1). In both the SIO and LIO simulations, however, far more free energy is dissipated by passive light absorption than it is by respiration (Fig. 12), although the difference is more striking in the SIO simulation (Fig. 12a versus b). Of course, some prokaryotes in nature harness the abundant light energy via expression of proteorhodopsin (Beja et al., 2000; Giovannoni, 2017).

**Fig. 12.**
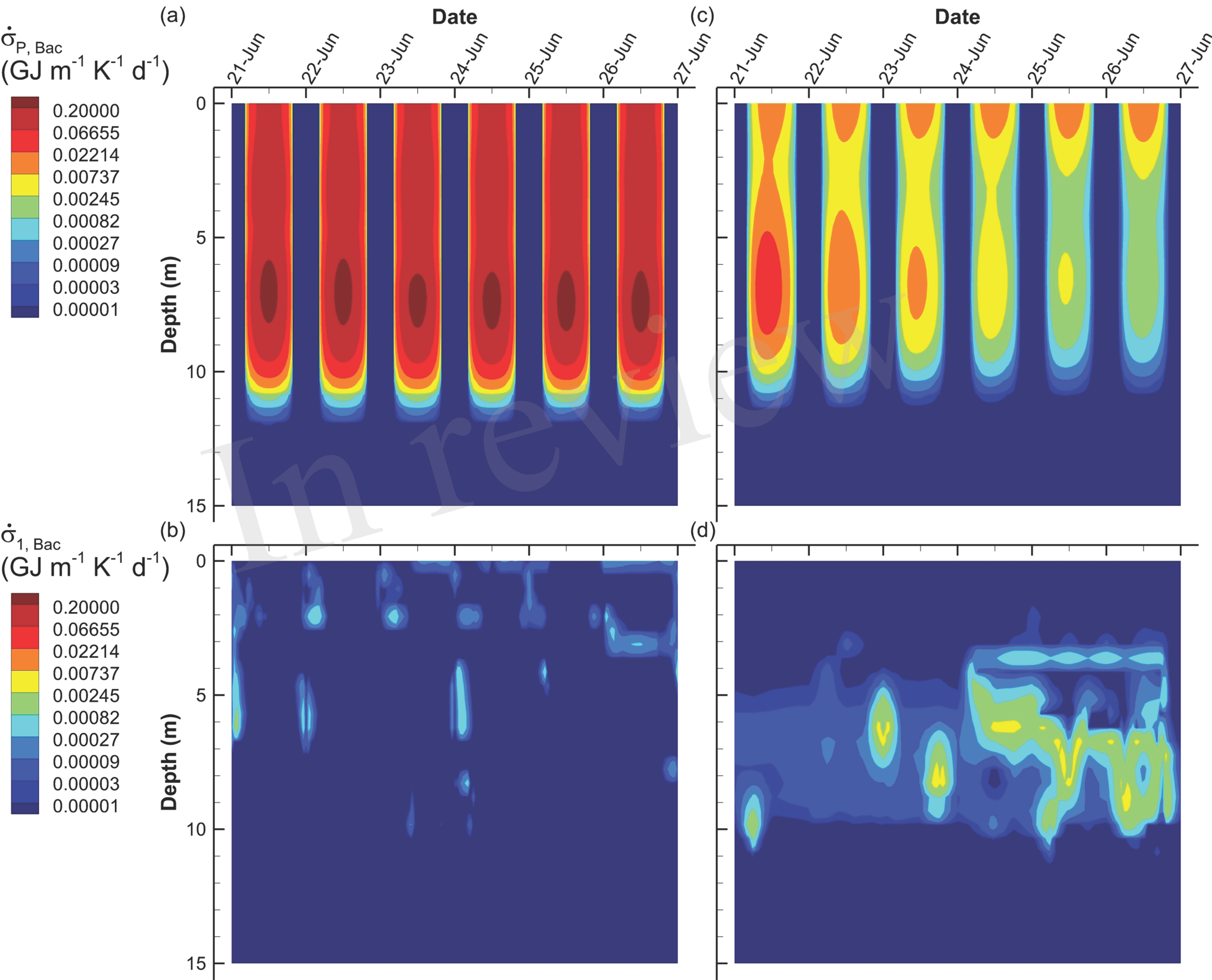
Contour plots of entropy production by bacteria associated with light absorption by bacterial biomass, 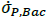 (a, c) and growth of bacteria, *r_1,Bac_*, on organic carbon, 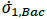, Eq. (S94) (b, d) for the SIO (a, b) and LIO (c, d) simulations.

## 4 Discussion

The two primary objectives of this modelling study were to demonstrate that 1) a model based on free energy dissipation reasonably describes microbial community organization and function and 2) that microbial systems operate collectively over characteristic timescales that are likely longer than what common wisdom would suggest. The secondary objectives were to demonstrate how the model can be used in systems with spatial dimensions and to extend the approach to include phototrophs. While improvements could be made with explicit data assimilation (Edwards et al., 2015), the MEP model did a reasonable job at simulating biogeochemistry in Siders Pond, and the better fit of the long interval optimization (LIO) simulation to observations indicates that the microbial community has evolved to function over time scales that are longer than 0.25 days. While sinking velocities and the two light absorption coefficients were crudely tuned (Section 2.1.3.1), the model otherwise has very few adjustable parameters compared to conventional biogeochemical models that often contain dozens (Ward et al., 2010). Furthermore, the MEP model is expected be more robust to extrapolation beyond the envelope of calibration data, as the MEP objective is applicable regardless of a systems operating point. The main drawback to the MEP approach is in the prediction of details. Since the MEP conjecture is based on statistical mechanics (Dewar, 2003; Lorenz, 2003), it does not provide information on the mechanisms by which a system attains an MEP state. If fact, the reason systems tend to follow an MEP trajectory is because there are many ways to achieve that objective. The approach is unlikely to predict details correctly. For instance, MEP models are unlikely to distinguish an algal bloom from a harmful algal bloom, although predicting the latter is of great importance. Likewise, an MEP model is just as likely to predict biomass turnover by jellyfish versus turnover by salmon, but from an economics perspective, these are two vastly different solutions. In essence, MEP is best used at predicting overall function, not the details that describe that function, largely because predicting details is very difficult and prediction accuracy decays rapidly. MEP modeling is similar to weather versus climate prediction. Climate models can predict far into the future, but not at the details of weather, and weather models provide details, but their prediction decays quickly. Perhaps the most useful aspect is that MEP provides a different perspective to view biology (Skene, 2017). For the Siders Pond model, the perspective was microbial functional activity over time and space.

Temporal strategies, such as circadian clocks (Wolf and Arkin, 2003), anticipatory control (Mitchell et al., 2009; Katz and Springer, 2016), energy and resource storage (Schulz et al., 1999; Grover, 2011), and dormancy (Lewis, 2010) are hallmarks of biology, yet they are often not given much consideration when theory and models are developed for understanding biogeochemistry, even though temporal strategies have also been observed in microbial communities (Ottesen et al., 2014). Here our results demonstrate that different organizational timescales dramatically impact biogeochemistry and how microbial communities function. For instance, the SIO simulation does not invest resources in phytoplankton growth, because over the short 0.25 d optimization, water dissipates more electromagnetic potential than a small increase in phytoplankton biomass over the short interval. Instead, the SIO solution allocates resources to green sulfur bacteria (GSB) and photoheterotrophs (PH) growth, possibly because sinking losses near the surface of the pond are greater than they are at depth due to advection and the pond’s geometry. Near the surface of the pond, upward advective velocity that counteracts sinking is only 0.01 m d^-1^, while at 12 m velocities reach 0.06 m d^-1^, which facilitates biomass accumulation. Specific growth rates need to be higher at the surface. The SIO solution also places more resources on decomposing refractory carbon, but also on respiring the produced labile carbon, which results in low standing-stock concentrations of C_*L*_. Grazing rates, especially under aerobic conditions, are higher in the SIO simulations as well. These types of resource allocations in the SIO solution are consistent with R* or resource-ratio theory (Tilman, 1982) and r-selection (Pianka, 1970; Fierer et al., 2007); that is, emphasis on fast growth. On the contrary, the LIO solution appears more similar to K-selection where resources are invested for longer term outcomes. In the SIO simulations, the control variables show rapid bang-bang control fluctuations as resources vary (Figs. 8d, S5a and S7a), while the LIO solution produces more gradual changes in control variables (Figs. 8i, S5d and S7d). Long-term strategies are more likely to produce cooperation that require investment of resources and more stable conditions (Hibbing et al., 2010), but utilize and dissipate resources more effectively (Cole et al., 2015). Strategies such as cross-feeding that make communities more resilient (Zelezniak et al., 2015) are also consistent with LIO solutions. A microbial community that implements temporal strategies should outcompete a community which lacks such temporal strategies. In essence, long-term strategies result in greater acquisition and utilization of food resources under non steady-state conditions than short-term strategies.

One of the main questions that arises in applying MEP is what is the appropriate timescale over which biology organizes? That is, in the current implementation of the model, what is an appropriate value for ∆*t* used in the optimal control problem (Eqs. (S183))? In this study two values were explored, 0.25 d for the SIO and 3 d for the LIO, but these were largely ad hoc choices. Our results indicate that the SIO solution does not match observations or expectations as well as the LIO solution. Comparing MEP model output to observations for differing values of ∆*t* is one means to explore this fundamental propery of ecosystems. But is a 3 d optimization window sufficient? It seems the appropriate MEP time scales do not depend on just the characteristic time scales of organisms, because our study on periodically forced methanotroph communities showed that the communities were well adapted to 20 day cyclic inputs of energy, even though the characteristic turnover time of bacteria is closer to hours (Vallino et al., 2014; Fernandez-Gonzalez et al., 2016). We know that terrestrial systems function on long timescales, at least on the order of the four seasons, but their time scales are likely much longer given that individual trees can live hundreds years, and forest succession takes longer than that (Odum, 1969; Finegan, 1984). At this time, we do not have an answer to the question regarding an appropriate optimization time scale, but it is likely related to the time scales over which biological predictions, acquired by evolution, can be reliably made, such as the sun will return tomorrow, or winter is coming. Predicting the future has obvious evolutionary advantages, but it also results in greater dissipation of free energy and entropy production. Some specific temporal predictions can be quite long, such as periodical cicadas that have strategies that operate on 13 and 17 year cycles, which clearly have been successful to date. We believe that determining the temporal scales that microbial communities operate over will be important for developing predictive biogeochemistry models, but time is not the only scale of importance.

Strategies that coordinate function over space also lead to greater free energy dissipation (Vallino, 2011). Some examples of such coordination include quorum sensing (Goo et al., 2012; Hmelo, 2017) and associated quorum policing (Whiteley et al., 2017), long-range metabolic signaling (Liu et al., 2015), stigmergy (Gloag et al., 2013), horizontal gene transfer (Treangen and Rocha, 2011), cables and nanowires (Reguera et al., 2005; Schauer et al., 2014), cross-feeding (Estrela et al., 2012; Rakoff-Nahoum et al., 2016), chemotaxis (Stocker and Seymour, 2012), vertical migration (Inoue and Iseri, 2012) and other types infochemical exchange (Moran, 2015). Our original intent was also to compare local versus global MEP optimization, but global optimization in the Siders Pond model did not produce biologically relevant results due to the speed at which electromagnetic potential is dissipated abiotically. Because water rapidly absorbs light, any water column of sufficient depth will dissipate all incoming radiation as entropy regardless of the presence of particles, so an infinite number of solutions exist that all produce the same global entropy production. For instance, a water column devoid of particles will produce entropy at an exponentially decreasing rate with depth, but eventually all the light will be dissipated. A solution in which the surface is particle rich will result in rapid entropy production with depth, but the same total amount of entropy will be produced. Particles placed lower in the water column would produce a different profile of entropy production, but the result would be the same, with all of the electromagnetic potential destroyed. Consequently, maximizing entropy production summed over the water column (i.e., global entropy maximization) does not produce a unique solution as it does for systems where energy potentials are stable, such as chemical potentials (Vallino, 2011). However, maximizing local entropy does select for a unique solution, which our results show is biologically relevant. In local maximization of entropy production, the objective is to maximize entropy production, 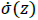, at every depth *z* simultaneously. The order in which 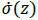 is maximized at each grid point does not matter, but in theory the optimization should be repeated until no further improvement occurs at any depth. Since light intensity is greatest at the surface, entropy production there can be greatly increased with an increase in particle density, which then lowers the amount of entropy that can be produced at depth due to shading. Of course, if the particles are phytoplankton and other organisms, resource availability will often limit biomass accumulation and local entropy production, but the system will organize in an attempt to minimize resource limitations, such as grazing and remineralization, and this is the solution observed in both the SIO and LIO simulations, with the latter outperforming the former.

This first MEP study involving light dissipation and phototrophy generated an additional interesting question--under what circumstances should entropy production be maximized locally versus globally? Our previous study with purely chemical reactions showed that global entropy production results in greater chemical potential destruction (Vallino, 2011). This study indicates that if abiotic processes can destroy an energy potential faster than biotic ones, then local MEP optimization will be the preferred solution. Global maximization of entropy production is more likely to be found in systems where energy potentials are stable with respect to abiotic decay, such as chemical potentials. However, it seems possible that both local and global optimization could be operating simultaneously is a system given that both light and chemical potentials exist. If so, one possible means of finding the correct solution is to first conduct local optimization followed by global optimization. In the case of the Siders Pond MEP model, a solution could first be determined from local optimization as was described in the Results Section, then that solution could be used as an initial condition for global optimization. If the global solution cannot improve on the local optimization, then the local optimization is the final solution. However, if a solution found by local optimization can be improved upon by a global optimization then that solution should be chosen. We did not conduct this study, as the MEP modeling approach is computationally demanding, and other components of the approach could use further development, which we briefly explain next.

The adaptive Monod equation that servers as the kinetic driver, *F_T_* (e.g., Eq. (S45) and others) is based on simple observations of bacterial growth under oligotrophic to copiotrophic conditions, but there must be underlying biophysical constraints that form the basis of the tradeoff between growing fast versus taking up nutrients at low concentrations (Kirchman, 2016). While the equation might be improved by using allometric relationships (West and Brown, 2004), determining the true limitations using genome-scale models may produce more robust tradeoff models for growth kinetics (Casey et al., 2016; Levering et al., 2016). Condensing genome-scale models may also be a means to construct more realistic whole community metabolic networks from genomic surveys (Hanson et al., 2014; Hanemaaijer et al., 2015), and exometabolomics could be used to identify metabolite nodes in the distributed metabolic network that are widely exchanged between functional groups (Klitgord and Segre, 2010; Baran et al., 2015; Fiore et al., 2015; Ponomarova and Patil, 2015). The reactions for predation in the Siders Pond model (*r_i,Gz_* and *r_i,AGz_*, Table 1) were designed so as not to introduce any new parameters (Vallino et al., 2014), but these reaction, which are intended to approximate virus to protist predation, is likely too simplistic and could use further development. Our reactions for carbon dioxide fixation (*r_1,Phy_*, *r_1,GSB_* and *r_1,SOx_*) could likely be improved by basing them on CO_2_ fixation pathways that have been discovered to date, which also reveals connections to their thermodynamic efficiencies, *ε_j_* (Bar-Even et al., 2012). Similarly, photoheterotrophy may be more widespread than appreciated, and there is a diversity of pathways (Kirchman and Hanson, 2013; Dubinsky et al., 2017), so our first attempt at representing photoheterotrophy could likely use refinement. Finally, the current approach uses non-linear optimal control to locate MEP solutions, but this is a computationally intensive problem, and falls under the class of problems known as control of partial differential equations (Coron, 2007). Building on expertise from that field could reduce computation requirements, especially for problems involving two and three dimensions, but it might be possible to avoid the formal optimization problem altogether. One possibility would be to use Darwinian inspired trait-based modeling approaches (Follows et al., 2007; Coles et al., 2017), which is, after all, how biology actually solves the MEP problem. The challenge in trait-based models is getting trade-offs correct, such as growth rate versus light level or resource storage. In this case, entropy production can be used to evaluate and cull models. For instance, when multiple versions of a model have been proposed, those models that produce more integrated entropy production over time and space should be selected for. MEP would also be useful for developing trait models that improve entropy production over time and space, such as resource storage, time delays (i.e., dormancy), migration across fronts and boundaries to acquire resources, clocks and oscillators, distribution and packaging of metabolic function, predation, remineralization and other regulatory motifs (Wolf and Arkin, 2003). By placing emphasis on the mechanisms of how energy potentials are destroyed over time and space, rather than on the peculiarities of how organisms grow and survive, can lead to new insights that can improve our understanding of biogeochemistry and model predictions thereof.

## 5 Conclusions

Development of the maximum entropy production (MEP) model is driven by the hypothesis that incorporating a fundamental principle in microbial biogeochemistry models will result in more robust extrapolation of model predictions. Our results demonstrate that models based on MEP can reasonably simulate how microbial communities organize and function in Siders Pond over time and space while using a minimum of adjustable parameters. The improved qualitative and quantitative agreement between model predictions and observations using long (3 d, LIO) versus short (0.25 d, SIO) interval optimization supports the hypothesis that biological systems maximize entropy production over long timescales. The modeling presented here extends the MEP approach to include an explicit spatial dimension, and new metabolic reactions were introduced to model phototrophs and entropy production associated with the destruction of electromagnetic potential. By considering the dissipation of both chemical and electromagnetic potentials, the MEP model shows that heterotrophs, such as bacteria, dissipate far more free energy in the photic zone by passive light absorption than by chemical respiration. Short interval optimization (SIO, 0.25 d) results in higher grazing rates and turnover of organic carbon, as well as rapid (bang-bang) changes in the reaction efficiency control variables, while long interval optimization supports higher phytoplankton growth and standing stocks near the surface of the pond. We also found that maximization of local entropy production, as opposed to global entropy production, must be used for energy potentials that are quickly dissipated by abiotic processes, such as light absorption by water and particles. However, our general conjecture that biological systems evolve and organize to maximize entropy production over the greatest possible spatial and temporal scales, while abiotic processes maximize instantaneous and local entropy production, remains valid.

## 7 Author Contributions

Both JV and JH contributed equally to project concepts and design, and both were full participants of the 24 hour endurance sampling of Siders Pond. Preliminary results from metatranscriptomics and metagenomics from JH were used to improve development of the metabolic network. JV developed the model and wrote first draft of the manuscript with editing and inputs during the process from JH. Both authors contributed to manuscript revision, read and approved the submitted version.

## 9 Acknowledgments

We thank Petra Byl for assistance in field work preparation and sampling of Siders Pond, as well as insightful conversations about the project. We also thank Jane Tucker for dissolved inorganic carbon analyses, Suzanne Thomas for hydrogen sulfide analyses, Sam Kelsey for dissolved organic carbon analyses, Kate Morkeski for sulfate, phosphate and particulate organic carbon analyses, and Rich McHorney, Leslie Murphy, Emily Reddington, and Gretta Serres for assistance with Siders Pond sampling logistics. We also thank the Semester in Environmental Science Program at MBL for use of their Hydrolab and Jon Boat use for sampling and a special thank you to Tom Gregg who allowed us to access Siders Pond through his property as well as for providing logistical support.

